# Traveling cortical netwaves compose a mindstream

**DOI:** 10.1101/705947

**Authors:** Ernst Rudolf M. Hülsmann

## Abstract

The brain creates a physical response out of signals in a cascade of streaming transformations. These transformations occur over networks, which have been described in anatomical, cyto-, myeloarchitectonic and functional research. The totality of these networks has been modelled and synthesised in phases across a continuous time-space-function axis, through ascending and descending hierarchical levels of association^1-3^ via changing coalitions of traveling netwaves^4-6^, where localised disorders might spread locally throughout the neighbouring tissues. This study quantified the model empirically with time-resolving functional magnetic resonance imaging of an imperative, visually-triggered, self-delayed, therefor double-event related response task. The resulting time series unfold in the range of slow cortical potentials the spatio-temporal integrity of a cortical pathway from the source of perception to the mouth of reaction in and out of known functional, anatomical and cytoarchitectonic networks. These pathways are consolidated in phase images described by a small vector matrix, which leads to massive simplification of cortical field theory and even to simple technical applications.

## INTRODUCTION

On a first sight it seems to be self-evident that a local perturbation within the cortical sheet should spread locally. The underpinning networks (see methods table II), dictating pathways, have been developed using different technologies and have varying numbers of nodes: anatomical^30-38^ cards are underpinned by fasciculi and smaller subcortical connectomes^48-52^ and structured by cytoarchitectonic^39-47,94-96,98-100,104,109,111,121-124^. These biological networks carry functions which, originally descriptively^54,55^ defined in psychological terms, lead, with integrated systems perspective^56,57^, to the low-dimensional multifocal cards from dynamic functional magnetic resonance imaging (dfMRI) seen in resting states^58,61,62,64-89^.

The networks are subject to a vectored dynamic system: in its multiple primary areas influxes of information flow inside and intermingle before running towards the only outlet in the primary motor area. This produces well defined temporal, spatial and functional boundary conditions. The system, even taking into account systemic feedforward-feedback-loops, yields an orientated direction and thus, using Hebbian^1^ principles as a basis, Mesulam^2,3,4^ develops a model (figure I) where a linear thought stream is represented via four levels of association. A hierarchical vector is created across the four levels, integrating the above-mentioned boundary conditions. This four-levelled phase space illustrates not only effective but also causal and structural connectivity by using orthogonal, inter- and intra-associative projections:

**Figure I.**
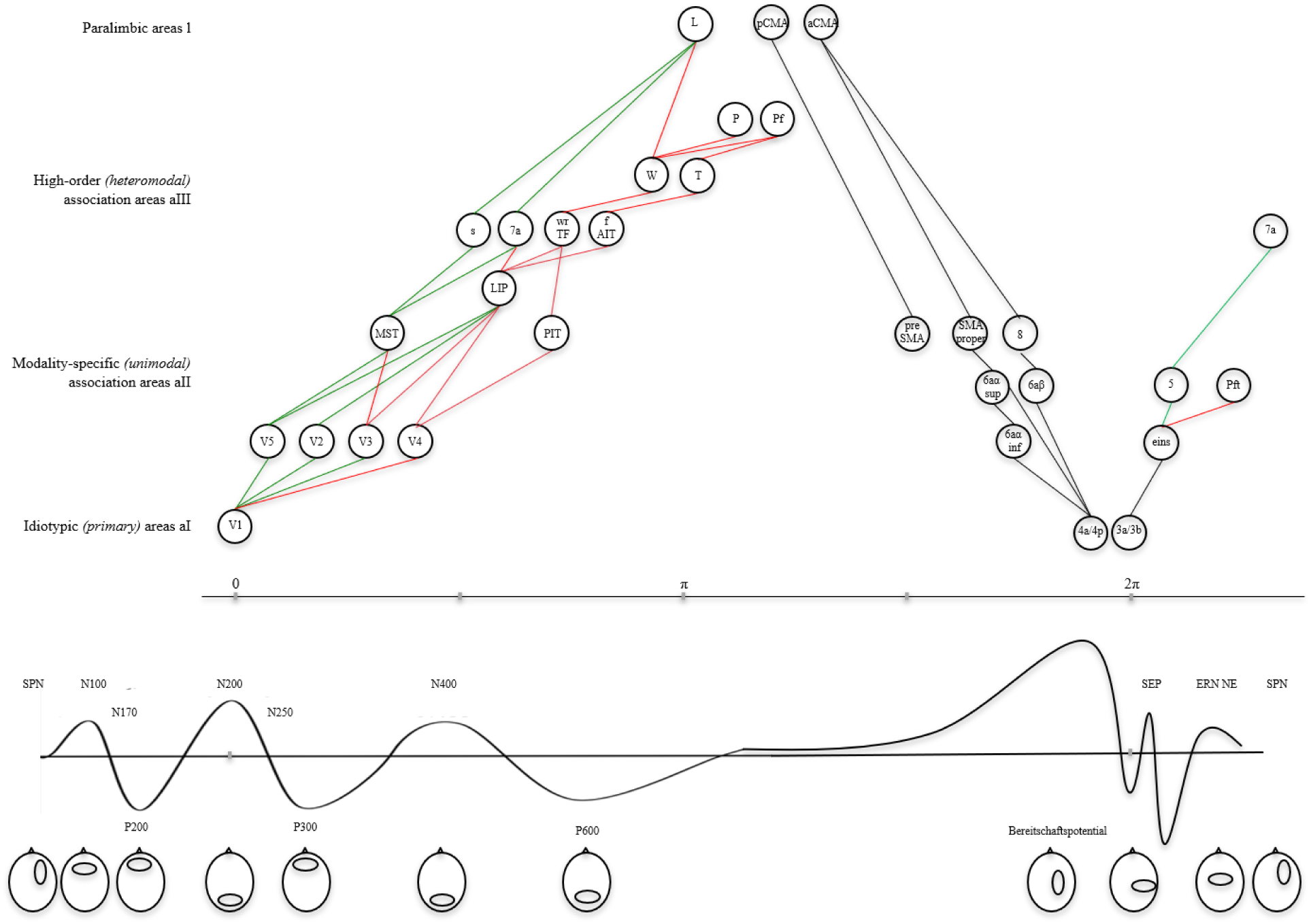
Hierarchies, longlasting waves and sensorimotor phases. Above - The phasic hierarchy model of a visuo-motor task, adapted from Mesulam^2^, figure 2, stretched over time. The nodes named (within circles) following Mesulam (on the left), Jubrain^45-47,63^ respectively (on the right), 2π from visuo-perception to auto-perception. AIT - anterior inferotemporal cortex; f - area specialized for face encoding; L - hippocampal–entorhinal or amygdaloid components of the limbic system; LIP - lateral intraparietal cortex; MST - medial superior temporal cortex; P - heteromodal posterior parietal cortex; Pf - lateral prefrontal cortex; s - area specialized for encoding spatial location; PIT - posterior inferotemporal cortex; T - heteromodal lateral temporal cortex; TF - part of medial inferotemporal cortex; V1 - primary visual cortex; V2, V3, V4, V5 - additional visual areas; W - Wernicke’s area; wr - area specialized for encoding word-forms; 7a - part of dorsal parieto-occipital cortex. Red lines – ventral streams; green lines - dorsal streams. Details see methods, figures IV-VII and movie I. Bottom - Event-related potentials ordered over time demonstrate like the rapid monosynaptic transmission-times in the range of 10 ms (from V1 to V5) to 50 ms (from V1 to AIT), as between Mesulam’s hierarchical levels, add up to longlasting waves in the order from 2 Hertz down to 0.25 Hertz. Note that Hebbian monosynaptic feedforward-backward loops occur in parallel to the slow frequency oscillations in the range of some 10 ms. SPN - Stimulus-preceding negativity, awaiting an informative stimulus; N100 – preattentive perceptual processing; N170 - faces; P200 – preattentive perceptual processing; N200 – stimulus detection; N250 – face identity; P300 – stimulus categorization and memory updating; N400 – semantic and conceptual processing; P600 – syntactic processing; SEP – Sensory evoked potential; ERN NE - Error negativity, false acts; Pe - Error positivity, false acts.

A cycling phase might be started with the perception of a stimulus in a primary, idiotypical, unimodal association area, as e.g. within the visual cortex (ai); discrimination occurs further upstream in a secondary, still modality-specific association area (aii); heteromodal and often conscious integration follows in a tertiary, high-order association area (aiii); and as highest association area, a limbic, emotional response will upspring (l). Further ahead in time, the chain of ai-aii-aiii in reverse, falls down in descending order via a tertiary multimodal level of association, as in prefrontal thought processes (aiii); the anticipatory planning of movements, as in the vast secondary unimodal, supplementary-motor area (aii); to the execution of the movement in the primary level of association (ai), the motor cortex – a process of execution which ultimately triggers the ai-aii-aiii chain once again: this time through self-perception which takes place in the primary somatosensory area (ai) over unimodal discrimination in secondary parietal regions (aii) to multimodal parietal regions (aiii), resulting in temporal models as shown in figure I and more detailed in the methods figures IV – VII and movie I.

Independent of complex, dispersed processes, the order of the sequence ai-aii-aiii-l-aiii-aii-ai should be retained like a palindrome as a quasi-periodic pattern and be continually repeated in parallel and with varying phase lengths, just as the perception-reaction sequence involved in systematic thought processes also remains chronological and recurs rapidly throughout the day and gradually over the course of a lifetime in an ever-repeating self-similar loop.

The model raises a series of prognoses regarding the where, what and when of a transformation whose content is ultimately arbitrary and of the likelihood of such a linear, successive combination occurring. Regulated by feedforward-feedback-loops, interferences appear over longer time and space periods as static or moving waves, described by Crick & Koch^5^ as

> *… a forward-traveling, … propagating net-wave of neuronal activity, (but is not the same as a wave in a continuous medium) … These net-waves represent at each point in time ‘coalitions’ … There are coalitions, which achieve winning status and hence produce [a] conscious experience … In the conscious mode, it seems likely that the flow is in both directions so that it resembles more of a standing net-wave*.

Deco & al.^6^ ground this dynamic into a neural field model where the above described neural masses rise out of spiking neurons, concluding *in summary, for any model of neuronal dynamics, specified as a stochastic differential equation, there is a deterministic linear equation that can be integrated to generate ensemble dynamics*.

The corresponding empirical spatio-temporal matrix ***M*** and its physical node parameters (such as location, volume, point in time, phase, frequency, amplitude) and edge parameters (such as distance, direction, duration, hierarchy, functions) are determined, using time-resolved functional magnetic resonance imaging (fMRI), both separately and as arrays in tasking, i.e. double-event-related brains, where a visual instruction was followed by a considered motor-reaction (see methods).

## RESULTS

The appearance of the resulting phase image (figure II, details supplementary figures VIII & IX, movie II) unfolds before the view a wave train, which reflects the flow of the visual stream, the dorsal and the ventral one, fulminant, and reproduces at first glance in great clarity the sensorimotor stream, ventral and dorsal stream well separated too.

**Figure II.**
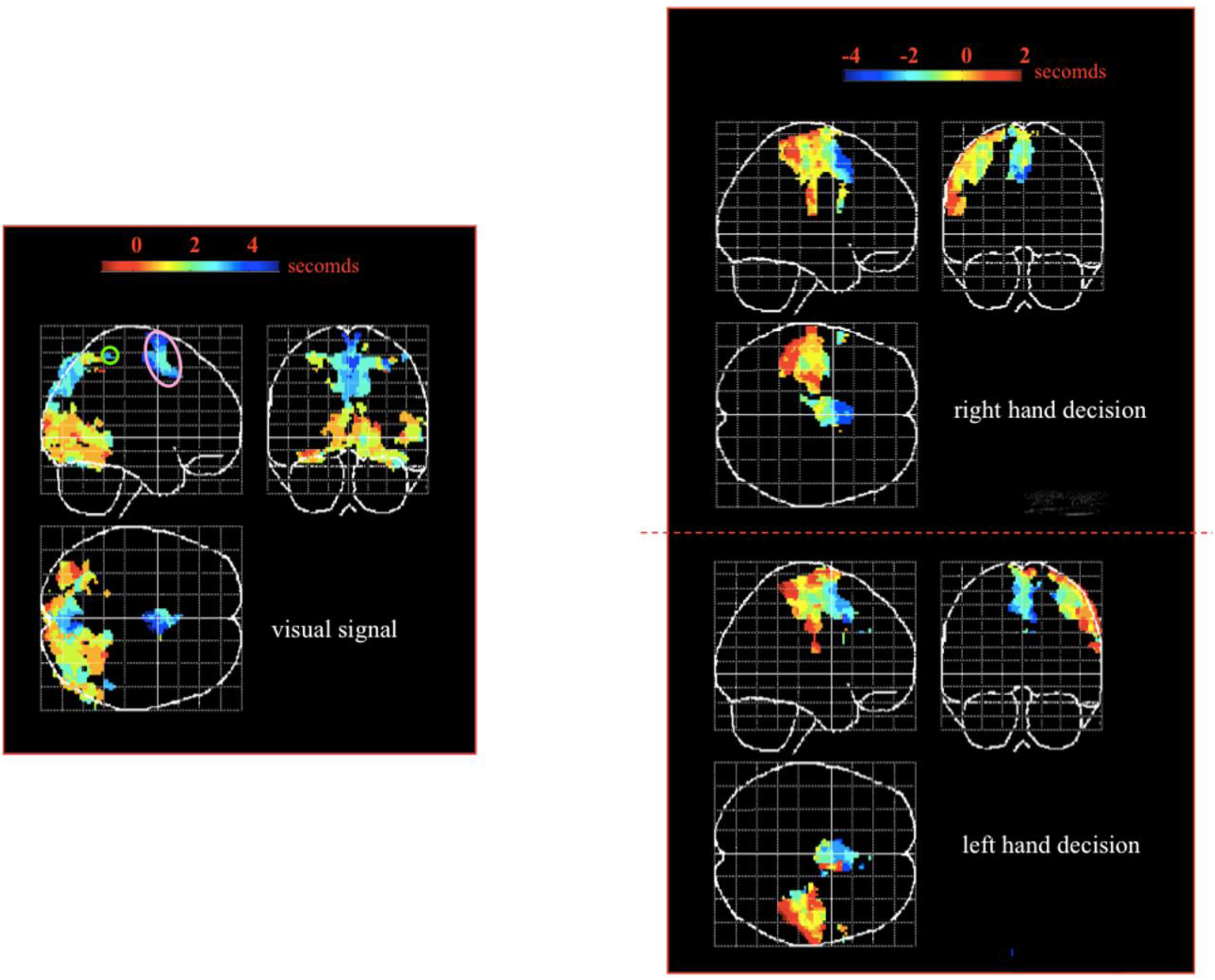
Phase images of a self-triggered response task. Phase images from time series (supplementary figures VIII and IX, movie II) show the dynamic of an optically triggered, imperative, self-delayed response task. Maximal-intensity-projections of the point-in-time of extreme activities as indicated by the colour bars. Left image event-related to the signal, and the two right pictures event-related to the respective reactions; stimulus-related from 1 second pre-stimulus (red) to 4 seconds post-stimulus (blue); reaction-related from 4 seconds prior to the reaction (blue) to 2 seconds after the reaction (red). The colour bar codes hierarchies^2,3,4^ too, separating unimodal (red/brown/green) from heteromodal (green/blue) areas. Note the spatio-temporal gradient spanning from the calcarine sulcus over the medial, lateral and inferior extrastriate cortex to the precuneus and superior parietal cortex, and from the deep mid-frontal pre-SMA to the upper SMA proper (purple circle), where a spatio-temporal gradient starts in the motor images, followed within the lateral convexity by gradients from the central sulcus to anterior and to posterior areas, 7PC/mip short cutting to the latter visual superior parietal side (green circle).

A right-left hemispheric symmetry of the activation pattern allows to limit the detailed analysis to one hemisphere, which will be the right one in the following detailed descriptions.

There is a medio-dorsal cluster in the visual phase image, the medial supplementary motor area SMA (figure II, pink oval), the latest cluster in time to have been activated. The same cluster is the first to be activated in the motor pictures through the movement. The two clusters in the two different phase images overlap in the right hemisphere over 356 mm^3^ (table I) and are constructed out of the same microanatomical, functional and temporal elements. Activation begins in the two conditions in pre-SMA (visual 3.1 s post, motor 2.1 s pre) and flows over the SMA proper to the medial upper border of the cortical mantle (visual 3.9 s post, motor 1.6 s pre).

**Table I.**
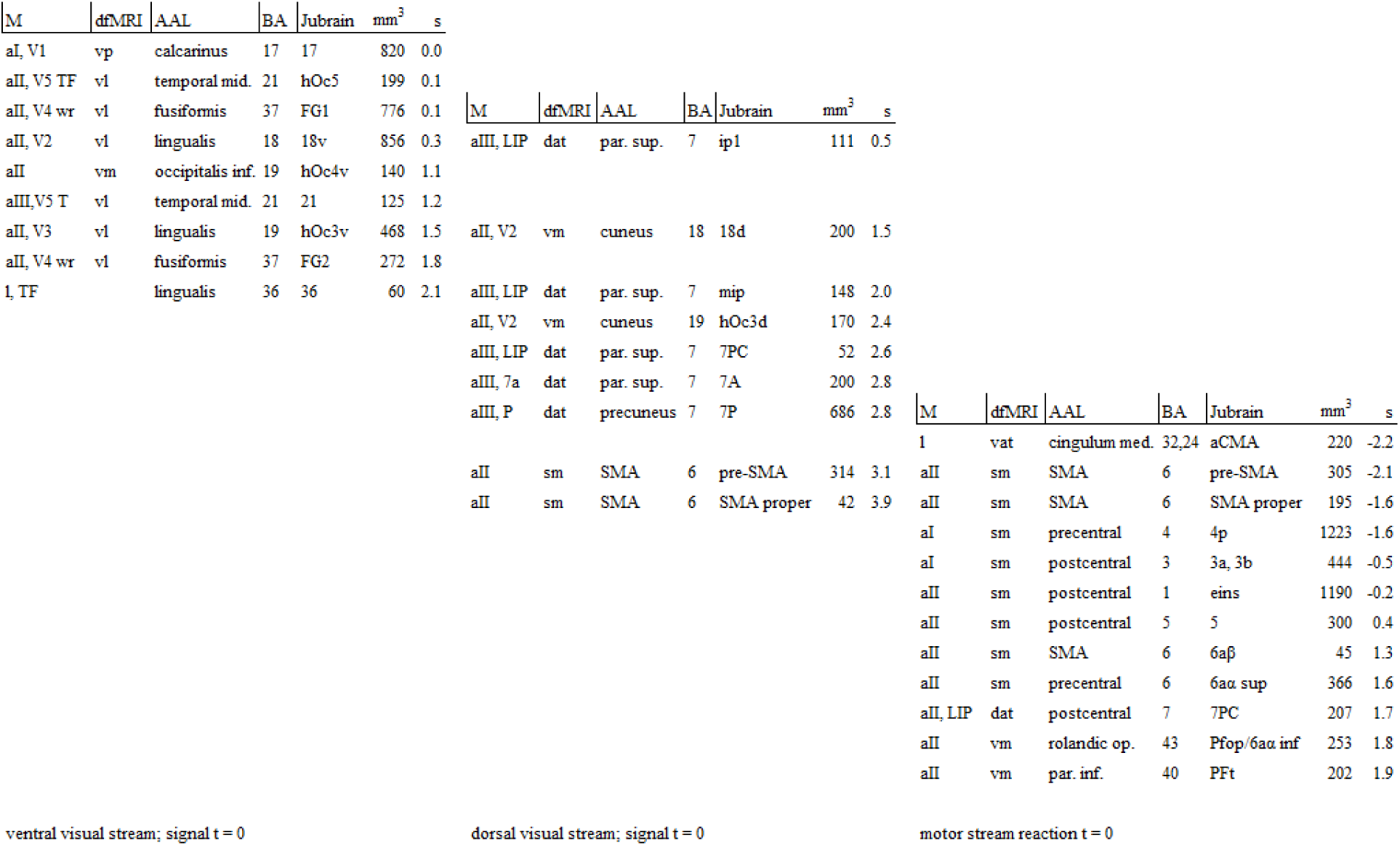
Hierarchical, functional, anatomical and histological visuo-motor nodes of the right hemisphere forming a mindstream. Columns: M - Mesulam^2,3^ (abbreviations as in figure I); dfMRI - dynamic functional resonance imaging (networks: vp – visual primary; vl – visual lateral; vm – visual medial; dat – dorsal attention; vat – ventral attention; sm – sensorimotor; vm – ventral motor); AAL, automated anatomical labeling^9^; BA – Brodmann^40,41^ areas; Jubrain^45-47^, but the differentiation within the large surface region 18 between 18v (18 ventral, gyrus lingualis) and 18d (18 dorsal, cuneus and gyrus occipitalis); also the as yet in JuBrain uninvestigated Brodmann areas 36 and 21; also aCMA, a small portion of BA 32 and the differentiation of the large surface volume of SMA according to myelographic classification^63^; mm^3^ – activated volume; s - the average points in time of maximal correlations for a given node in seconds related to the event.

A joint time period is therefore consolidated in which a time arrow of 5.2 s represents the time from one perception over cognition to action and back to autoperception, the phase 2π in Mesulam’s model out of figure I, with wave fronts covering expansive, anatomical, cyto- and myeloarchitectonical, functional and hierarchical areas, parallel and perpendicular, as shown in table I:

i) Anatomical network architecture is quantified (table I, column 3; both hemispheres supplementary table II) with Automated anatomical labeling (AAL)^32^ over 15 out of the 44 AAL-nodes (calcarine, temporal, fusiform, lingual, occipital inferior, parietal inferior, cuneus, parietal superior, supplementary motor area, middle cingulum, precentral, postcentral, Rolandic operculum and once more parietal inferior in each hemisphere). The AAL-algorithm gives activated volumes accumulating in total to almost 10000 mm^3^ in each hemisphere; with an outer layer thickness^7^ of 2.3 - 2.8 mm a neural coverage of about 250 cm^2^ for a surface^8^ of 1200 cm^2^. Identifying first, extreme and last point in time of correlation allows modelling of the regional *standing netwaves*, the frequencies moving with 0.3 – 0.5 Hz in the lower delta band. The entire set of *standing netwaves* results in a stream of continuously phase-shifted waves, where the individual regions develop in parallel as *winning coalitions* and form a *forward traveling netwave* across time and space.

ii) The cytoarchitectonic network (table I, columns 4 - 7) resolution augments the number of nodes up to 26 out of the 64 nodes shown in figure V. The ventral visual stream is here composed of 17^94^, FG1^91^, ventral 18^94^, hOc4v^95^, hOc3v, FG2^91,96^, BA 36, with a midtemporal branch hOc5^99^, BA 21^97,100^. The dorsal visual stream is composed of 17, dorsal 18, hOc3d^98^, ip1^109^, mip^104^, 7PC, 7A, and 7P^105-108,124^ and finally, a new observation, BA 6, pre-SMA ^63,110-116^.

The corresponding volitional motor response is made of a medial part from paleocortical aCMA^113^ over pre-SMA, SMA proper to 6aβ; and a lateral part from 4p^121^, a dorsal stream to prefrontal 6aα and to postcentral 3, to 1^122,123^ and further on to 7P^124^, as well as a ventral stream to 6aα inferior^63^ and to caudal PFt, Pfop^100^.

iii) These macro- and microanatomical architectures port dfMRI-correlations (table I, column 2). Their temporal architecture is obvious in a sequence over seven networks with sixteen nodes, representing their hypothesized functional succession: At first, consecutive visual networks^62,64,66,69,71,78-80^ turn up - the occipital, lateral and medial visual network as per Smith & al.^64^, with the latter two forming the high-level visual network, as per Zhang & al.^78^, including the six hierarchical nodes of Yeo’s & al.^62^ visual hierarchies.

Within the higher association areas it is the dorsal attention network^62,70,72,75,78-81^, which follows parietal and ultimately the medial-precentral part of the sensory motor network^64,65,78-81^ originates with the latter.

This and the anterior cingulate cortex, a part of the ventral attention/salience network^62,69,72,78,79^, is also the origin of the motor branch which subsequently employs the ventral motor network^78^, and flows posteriorly and anteriorly back into the dorsal attention network, and inferiorly towards the Roland’s motor operculum in the direction of the insular lobe, spreading in an almost elliptical fashion from the sulcus centralis.

iv) The hierarchical architecture (table I, column 1), including dynamic ideas, is of particular importance and shows the phenomenon of a linear *traveling netwave* over the visual perception, discrimination and higher association levels as delineated by Mesulam^2,3^. This demonstrates the starting point in early idiotypical, primary areas and its evolution over time via secondary unimodal to multimodal high association areas, located midparietal. These midparietal high-association areas are omitted in a speeded response task, demonstrating their high hierarchical position (supplementary figure X).

The reversed hierarchy is present in the movement-linked development from limbic regions over secondary to primary unimodal motor, to primary, secondary unimodal somatosensory and tertiary heteromodal sensory areas. Over the course of time the process recruits all the four Mesulam hierarchies over nine nodes.

The complex concordance from i) - iv) is grouped into large clusters, which radically simplifies the description to the determination of local and temporal extreme values (figure III). Vectors (black in figure III) are constructed between these extremes over the extension of those clusters, which in their orientation cover the above described, ordered sets of anatomies, cytoarchitectures, functions and hierarchies. Putting it together leads to a vector matrix bridging the combined visual occipito-temporo-parietal-midprefrontal, visual occipito-lingual-fusiform and sensorimotor prefrontal-precentral-postcentral-parietal networks. The matrix spans a field from the source of the visual perception through the conscious reflexion to the action output into the newly emerging autoperception. By way of the constructed vectors, a wave develops like Hayne’s *stream of consciousness*^9^.

**Figure III.**
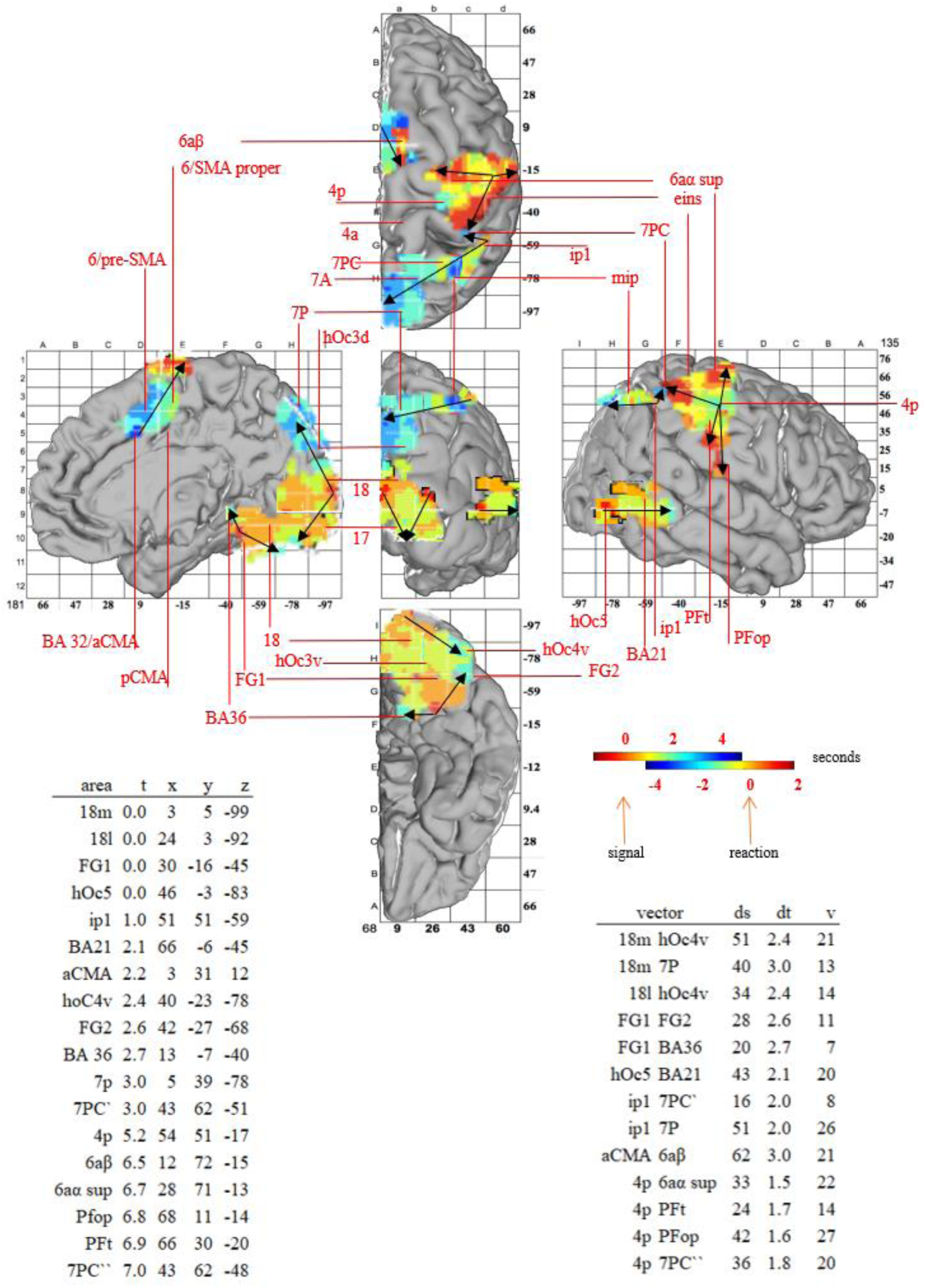
Dynamic topology of a visuo-motor vector field. The right hemispheres from figure II combined to an entire stream, surface-rendered and the histological architecture as per the definitions of JuBrain^45-47^, with the exception of BA 21 and 36, which are not yet published. The subclassification of SMA/BA 6 as per Zilles & al.^63^. 18m – medial (cuneus), 18l - lateral part (gyrus occipitalis) of dorsal JuBrain-area 18. The colour bars provide the time scale in reference to visual perception and movement in seconds (supplementary movie III). Temporal extreme values were determined within the regions and vectors extrapolated (black arrows). The table to the left provides the temporal extreme values (t in seconds, the time-arrow of the reaction transposed by -5.2 seconds to visual perception) and their point in space (x, y, z – Talairach^28^-coordinates in mm). The table to the right provides the vectors with their Euclidian distances in time (dt, in seconds) and space (ds, in mm).

Thus, the wanted matrix ***M*** is well approximated.

## DISCUSSION

Waves are built from their amplitude and their duration in space and time, oscillate in different frequencies, appear time- and space-locked to events, can spread locally *and are believed to be fundamental mechanisms of neuronal computation and interregional communication*^27^.

Neurons are the elemental building blocks of such cortical waves, spiking predominantly in the form of propagating, traveling waves, while at the meso- and macroscopic levels, mainly stationary waves have been described so far. Such stationary signals have been explored regarding the significance of their physical properties, as especially frequency. Slow cortical potentials, < 4 Hz, so in the time range of the here observed signals, have been seen by He & Raichle^10^ as optimally positioned for carrying out large-scale information integration in the brain, have even been linked by them to conscious processing, and have been inverted by neurofeedback^11^.

In fMRI the temporal properties of oscillations could be used to describe the standing, anticorrelated bumps on unimodal networks superficially and on multimodal networks very profoundly. Time-locked event-related standing oscillations are fundamental in electrophysiology (see examples of event-relateted potentials in figure I) and functional MRI, and even more information seems to be linked to phase-coded signals. Montemurro & al.^12^ calculate in macaques *that at low field potentials frequencies, the phase of firing conveyed 54% additional Shannon information beyond that conveyed by spike counts*. Glerean & al.^58^ demonstrate by using instantaneous phase synchronization as a measure of dynamic, time-varying functional connectivity in the visual cortex of man the same regions to be activated as in the present study but the mid-precentral SMA.

Lakatos & al.^13^ show even – again in the same wave-length as in the present study - that *stimulus-locked phase-coding in the delta frequency range are present when attended stimuli are in a rhythmic stream. These … oscillations … entrain to the rhythm of the stream, resulting in increased response gain for task-relevant events and decreased reaction times*.

However, traveling netwaves have only since recently, with the dawn of methods as multielectrode array, electro-corticography, voltage-sensitive dye, low field potential and intracellular recordings, been in the focus of interest. Traveling netwaves over some hundred’s milliseconds, observed in the cat’s visual cortex, voltage-sensitive dye registration in V1, V2 and V4 of the awake monkey, traveling beta oscillations during movement planning in macaques, and especially the electrocorticographic slow oscillations, 0.1 - 1.0 Hz, orchestrating large-scale neural spiking in human slow-wave sleep, traveling anterior-to-posterior with 1.2 - 7.0 m/s, lead Muller & al.^14^ to the conclusion *that stimulus-evoked travelling waves are a general and important feature of dynamics in the visual system …* so *… that dynamic wave fields introduce a general conceptual framework*.

All these individual biophysical properties - on standing waves, on frequencies, on rhythmic oscillations, on timelocking, on phase synchronization, on wandering waves – as observed in those studies are continuously and in overwhelming totality and simplicity present in the vector-matrix constructed in fig. III. The propagation speeds are from 0.1 to 0.25 m/s, the frequencies of 0.5 Hz, traveling across several centimeters, and adding up to a complete stream of about 50 cm length and 10 s duration. These waves develop parallel and dense in temporo-spatial continua, so that a current grows out of them. The phases of the current waves are temporally synchronized in a continuum along spatio-hierarchical-functional axes.

Methods to exploit the information encoded in time-varying connectivity have been developed in recent years in order to reach beyond the static measures of resting fMRI^15,16^. Task positive paradigms are, in their nature, even more proximately linked to time-varying phenomena. While resting state might resemble testing an orchestra preparing for the concert, a task positive brain would be like observing a playing orchestra, where time synchronization will occur as the result of supervised Hebbian learning. Finally, it is the task-positive space where we look for orbits and attractors^17^. However, the solution space shown here can now be used as seed-point series for dfMRI.

In the course of these considerations, the phases cluster were understood as Hebbian layers, which adhere to each other spatially. The synchronized stream that was observed here would, however, not represent the Hebb’s error minimization but it’s result. Thus high synchronization, sharing information between entities of a system, might be, far away from concrete, local functions as the perception of colour, intention or emotion, the essential Hebbian supervisor.

## CONCLUSION

Slow cortical potentials are ordered to a spatio-temporal matrix ***M*** out of a series of vectors, forming together a *stream of consciousness*^9^.

Well evaluated neuro-biological networks are reflected in ***M*** in a linear formation: The temporal order is reproduced i) between the dynamic functional networks visual, dorsal and ventral attention, and sensorimotor; ii) between the anatomical regions contained within these networks; iii) between the Brodmann and Zilles areas also found in these regions; iv) inside these areas and v) via functions and hierarchies.

Geometric waves are observed ‘hovering’ in the event of high over time synchronisation^18,19^ between input and output, in a similar fashion to a taut bow string. Crick & Koch’s^5^ concept of consciousness as *a forward-traveling, propagating net-wave of neuronal activity* holds true and its quantification as phase-shifted synchronisation^20^ includes the simplification of biophysical dynamic causal models^21,22^, as well as cortical field theory^6^.

The Mesulam and more deeply stacked hierarchical models can be linearly parametrised in task-positive networks accordingly.

The discussed models of Hebb^1^, Libet^114^, Kornhuber & Deecke^115^, Birbaumer^108^, Crick & Koch^5^, Mesulam^2,3^, He & Raichle^10^, Kjaer & al.^106^ and Haynes^9^ are essentially about the formalization of consciousness. It seems on the first sight evident that a complex system like consciousness should be composed out of even few and simple elements like other natural phenomena are constructed from simple elements. These elements, vectors, compose ***M***, which can now be inverted^23^ in a computational form leading, like Birbaumer & al.^108^ did in neurofeedback with standing slow cortical potentials, to dynamic applications as multi-channel transcranial stimulation, inducing a virtual reality – spectra of cortical cycling phase continua forming a mindstream.

## METHODS

The paradigm^24,25,26^ required a visual signal which prompted a test subject to push a button using either their left or right hand following a period they felt was “up to around 15 seconds”, with this process monitored using an fMRI scan:

Ten subjects (age range from 21 to 59 years, average age 30 years, 4 female, 6 male, all right-handers, as determined by the Edinburgh Inventory of Handedness) were studied. Informed written consents and approvals by the local ethic commission were obtained. The subjects held a button in each hand and watched visual stimuli presented from a video beamer onto a screen. The stimuli consisted of a short white flash (150 ms) on a black screen, followed by the appearance of a white arrow, which pointed in a pseudo-randomized sequence either to the right, or to the left. The arrow remained visible over the whole trial. The subjects were instructed to press the button with the corresponding thumb (left or right). They were furthermore instructed to choose, for each button press, varying temporal delays from zero up to approximately 15 seconds following the instruction flash. The whole experiment required 20 minutes, during which 87 trials of this delayed response task were presented with pseudo-randomized interstimulus-intervals of 14 +/- 2.5 seconds. The delayed task was contrasted with a speeded response task.

Event-related functional Magnetic Resonance Imaging was performed with a 1.5 Tesla MRI scanner (Siemens Vision, Erlangen, Germany). Twenty-four axial slices with 5 mm thickness and 3×3 mm^2^ in-plane resolution, 64^2^ matrix, 8/8 192 mm^2^ FOV covering the whole brain were acquired using a gradient recalled single shot echo-planar-imaging sequence. To shorten examination time TE was set to 40 ms. The measurement time for one whole brain acquisition was 2.32 seconds and volume repetition time was 2.52 seconds. Thus, for each experiment, 480 whole brain measurements were performed during a time interval of 20 minutes resulting in a mean of 5.5 volume measurements for each trial.

The image evaluation was performed with the Statistical Parametric Mapping programme package (SPM99, Wellcome Department of Cognitive Neurology, London). Preprocessing of the single subject time series images consisted of i) slice-timing, referring to the first slice (top), ii) realignment with sinc interpolation and adjusting for sampling errors, iii) non-linear normalisation to the canonical echo planar target image in Montreal Neurological Imaging space, interpolated with 2 x 2 x 2 mm^3^ voxel size, in order to achieve maximal spatial signal-resolution in the group statistics, and iv) spatial smoothing with a Gaussian filter with full-width half-maximum of 6 mm.

To generate time-resolved statistical maps, a series of models was calculated with time-shifted event delays. For each delay, the BOLD response was modelled using a generic hrf (global scaling, specified high-frequency-filter, using the hrf as low-pass-filter, which corresponds to a Gaussian filter with a cut-off level of 4 seconds full-width half-maximum, and without correcting for further internal correlation), resulting in separate t-maps for time points prior to the signal and the self-paced movements (−2 s for the signal, -3.5 s for the movements) and following (6 s for the signal, 2.5 s for movements). T-values were scaled for each volume proportionately from threshold (p < 0.001) to individual volumes extreme t-value at point in time t_e_. The resulting values were assigned to a baseline corrected^27^ activity I. Spatial segmentation used the algorithm of Tzourio-Mazoyer^32^, leaving only regions which excised 10% volume activity, except supplementary, figure X, insert i.

Table II shows cortical networks that served for comparison with the classification of the results.

**Table II.**
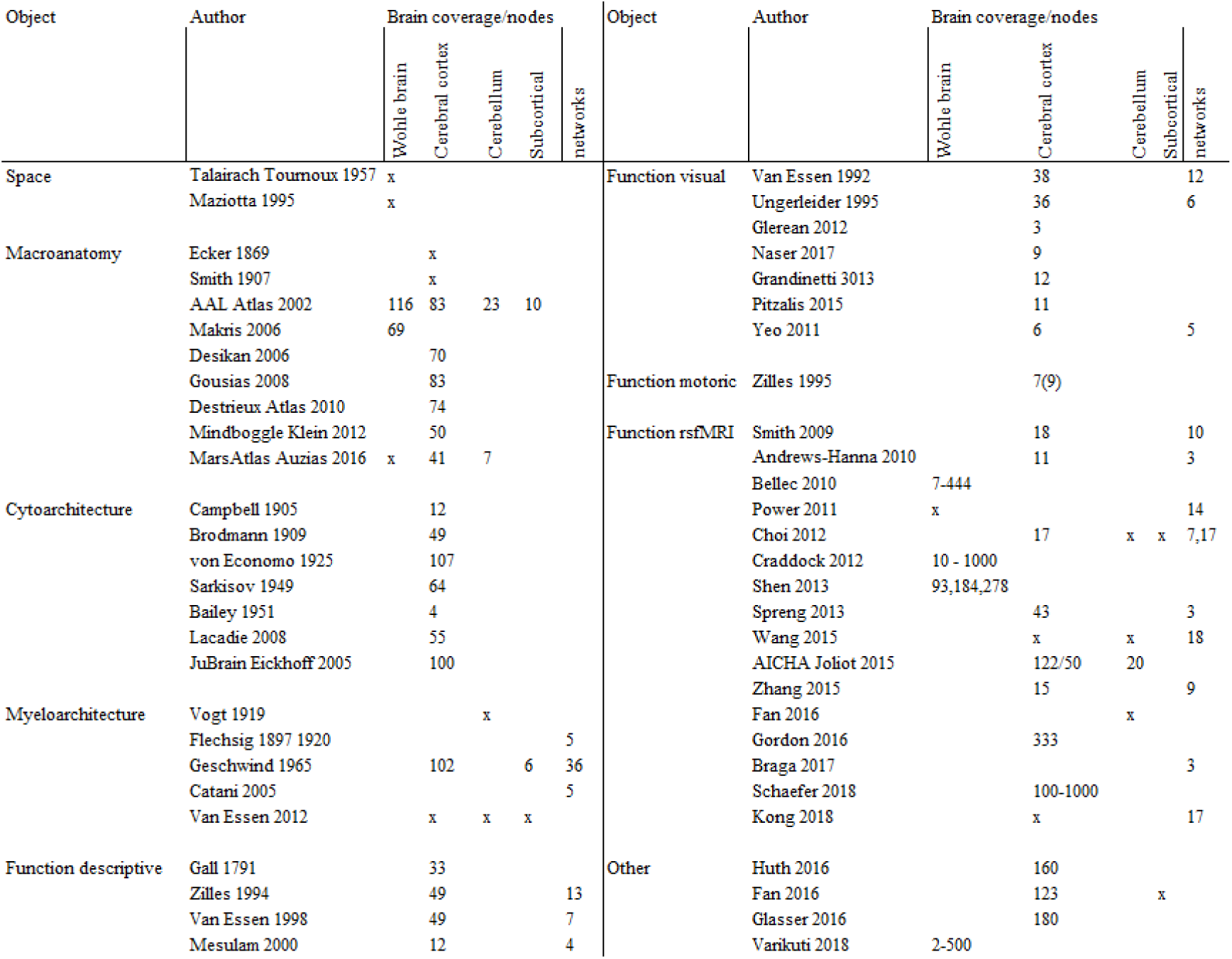
Nodes and networks in parcellations of the human brain. The space, the anatomy, the cytoarchitecture, the myeloarchitecture and the functions supported by the aforementioned structures, for the whole cortex as well as for the visual and motoric subsystems. Given are the nodes (per hemisphere) and, if investigated, their organization in networks and hierarchies. i) Space – constituent Talairach^28^ and MNI^29^ spaces. ii) Macroanatomy - Different anatomical subdivisions, as per old observations such as Ecker^30^, Smith^31^ and newer ones as Tzourio-Mazoyer & al.^32^, Harvard–Oxford Atlas Makris & al.^33^, Desikan–Killiany Atlas Desikan & al.^34^, Hammersmith Atlas Gousias & al.^35^, Destrieux & al.^36^, Klein & al.^37^, Auzias & al.^38^. iii) Cytoarchitecture – Campbell^39^, Brodmann^40^ and its transcription by the Yale Bioimage Suite Package Lacardie & al.^41^, von Economo & Koskinas^42^, Sarkisov & Filimonoff^43^, Bailey & von Bonin^44^, Zilles & al. ^45,46,47^ Torso. iv) Myeloarchitecture - as per Vogt & Vogt^48^, Flechsig^49^, Geschwind^50^, Catani & Ffytch^51^ and the ongoing efforts of the Human-Connectome-Project^52^ on cortical parcelling. v) Function descriptive – Gall ^53^, Zilles & Rehkaemper ^54^, Van Essen & al. ^55^, the two latter ones consolidating Brodmann’s atlas. vi) Function visual – Van Essen & al.^56^ (12 hierarchies), Ungerleider^57^ (6 hierarchies), Glerean & al.^58^ (dynamic regional phase synchrony), Naser & al.^59^, Grandinetti & al.^60^, Pitzalis & al.^61^, Yeo & al.^62^. vii) Function motoric - Zilles & al.^63^ (integrating architectonic, transmitter receptor, MRI and PET data). viii) Function dfMRI – resting state functional magnetic resonance imaging observing networks (default mode network (consisting of a core, a dorsal medial and a medial temporal subsystem), executive control network, frontoparietal network, dorsal and ventral attention network, salience network, several visual and sensorimotor networks): Smith & al.^64^ (ICA, 7000 subjects), Andrews-Hanna & al.^65,66^ (hierarchical relations within the default network), Bellec & al.^67^, Power & al.^68^, Yeo & al.^62^ (clustering ICA, 1000 subjects, separation of the early sensory and late motor cortices), Buckner & al.^69,70^ (seedbased ICA), Choi & al.^71,72^ (caudal-rostral parietal and rostro-caudal frontal hierarchies), Craddock & al.^73^ (spectral clustering), Shen & al.^74^, Spreng & al.^75^ (hierarchical relations between three networks), Wang & al.^76^ (interindividual), Joliot & al.^77^, Zhang & al.^78^ (directionality, subdividing two visual networks), Fan & al.^79^ (eigenentropy), Gordon & al.^80^ (individual), Braga & al.^81^ (within individual), Schaefer & al.^82^ (clustering and local boundary detection), Kong & al.^83^ (individual, character-predictive), ix) Other – Huth & al.^84, 85^ (language), Fan & al.^86^ (multimodal, structure, connectivity, function), Glasser & al.^87^ (multimodal), Varikuti & al.^88^. See Eickhoff & al.^89^ for a detailed discussion.

**Table III.**
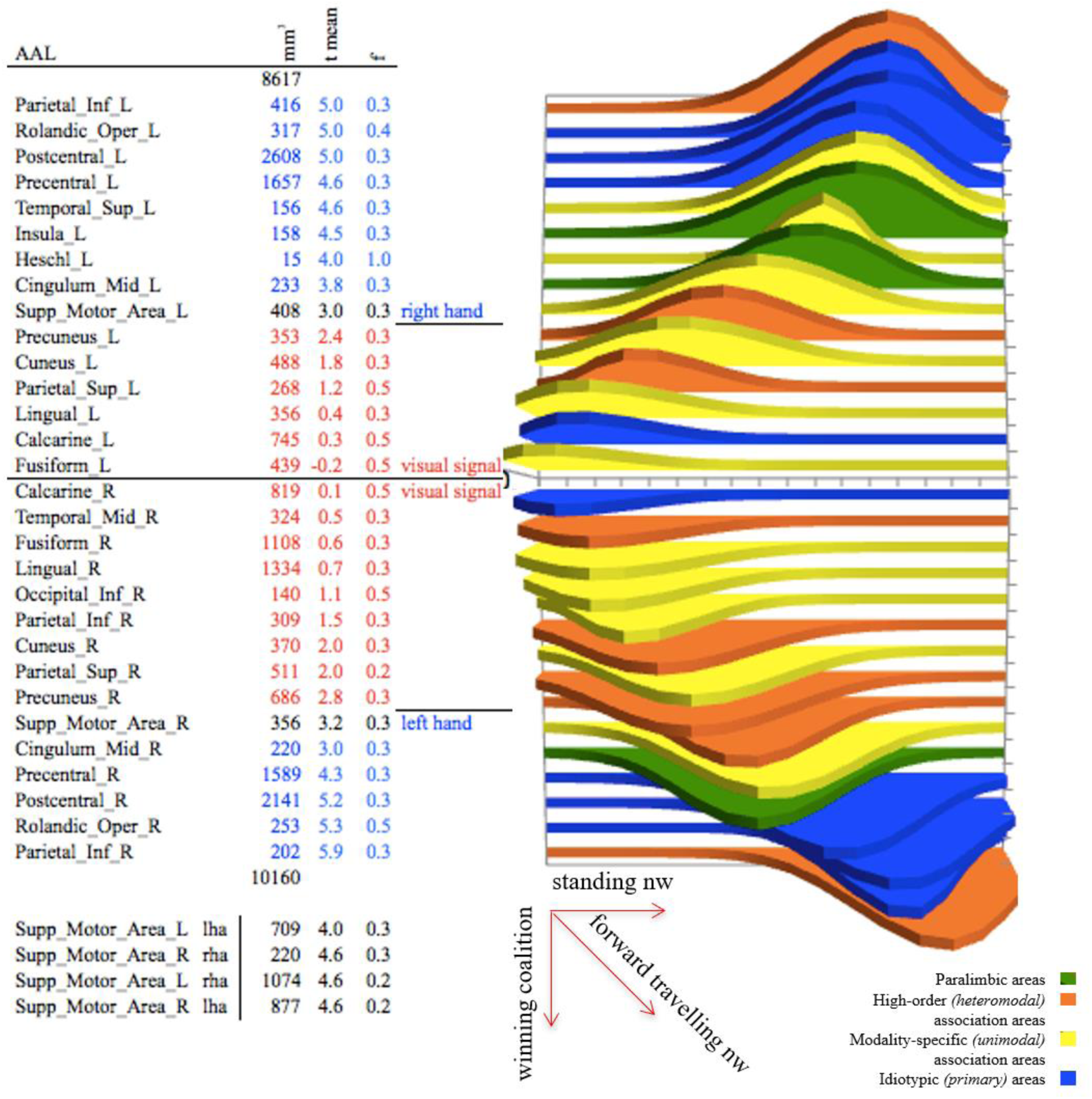
Right and left hemisphere in the AAL^32^-network. Visual signal, moving left and right hand (blue digits); AAL - Anatomical regions following Tzourio-Mazoyer^32^; mm^3^ – activated volume; tmean – mean-time of maximal activation over all voxels within the region in seconds, relating to the signal = 0 s (standard deviation 0.5 to 1 s); blue numbers for the movement, red for the signal. Black numbers indicate that this region activated under all conditions (the results for the SMA of the visual branch in the table, the results for the other conditions below); f – frequency in Hertz; lha – left hand; rha – right hand; nw – netwave. The colouration of the waves corresponds to Mesulam^2^. The individual *standing netwaves* are approximated by

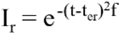

where *r* is the coordinate, its base-normalised regional nodes activity *Ir*, the time *t*, the mean time point *ter* of the maximum regional activities, and frequency *f*. Phase-shifted waves develop in parallel as *winning coalitions* and form a *forward traveling netwave* across time and space.

Figure IV shows the Jubrain-architecture^45-47^ aligned with the Mesulam model of hierarchies^2^, simplified in figure V, used to sculpture in figures VI and VII models of cortical streams that served for comparison with the classification of the results.

**Figure IV.**
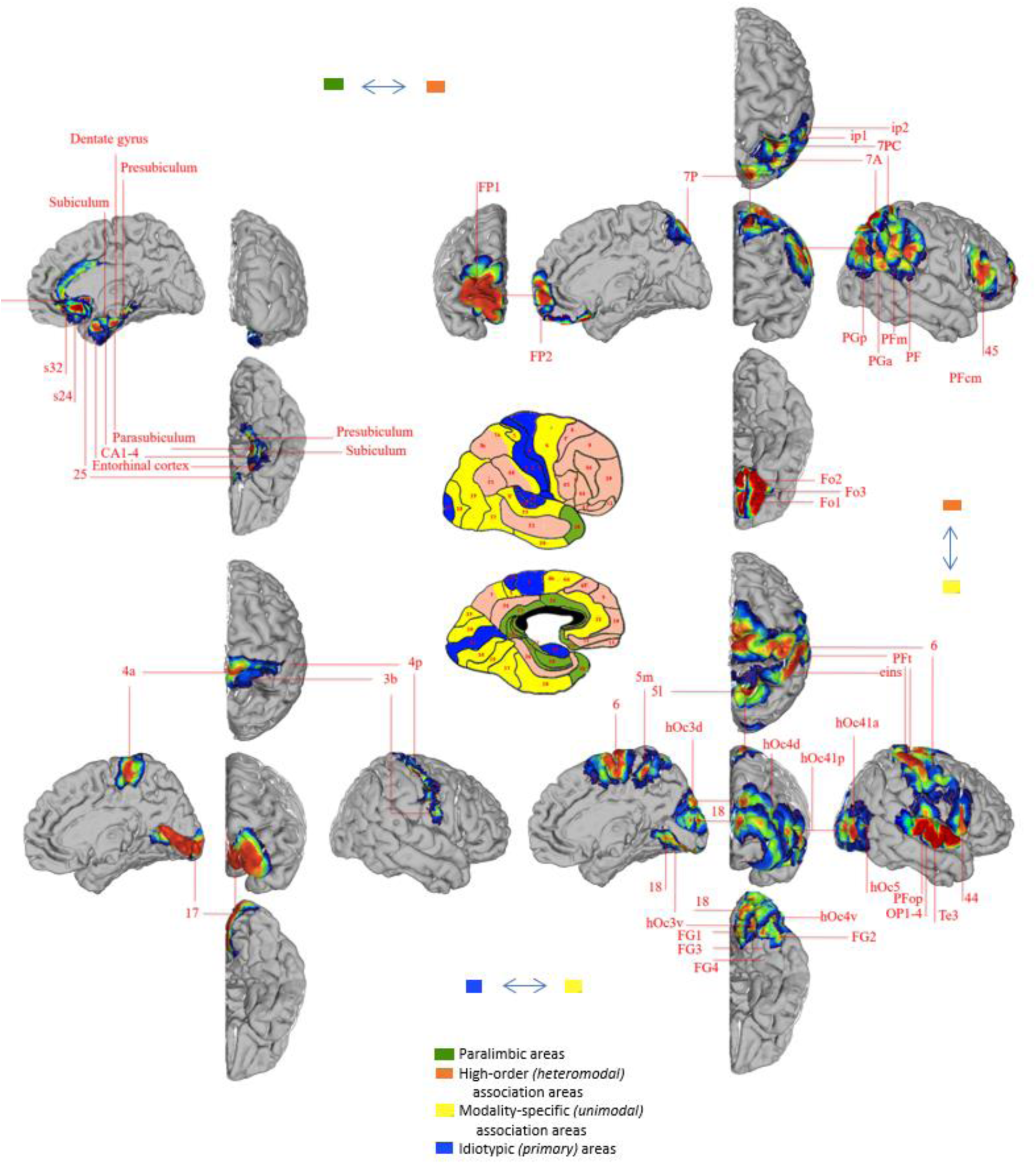
Temporal and spatial phasic hierarchies in Mesulam’s^2^ anatomical/histological/functional model (centre, adapted from Mesulam^3^) translate to the cytoarchitectures published in JuBrain^45-47^ (see methods movie I). The right hemisphere, unfolded. Waves might find their way in any direction, as, e.g., the top-down effect of the Bereitschaftspotential^115-116^ or the top-down effect of attention in early sensory cortex, largely invisible to spike recordings, but readily seen in the fMRI signal of monkeys^13,90^.

**Figure V.**
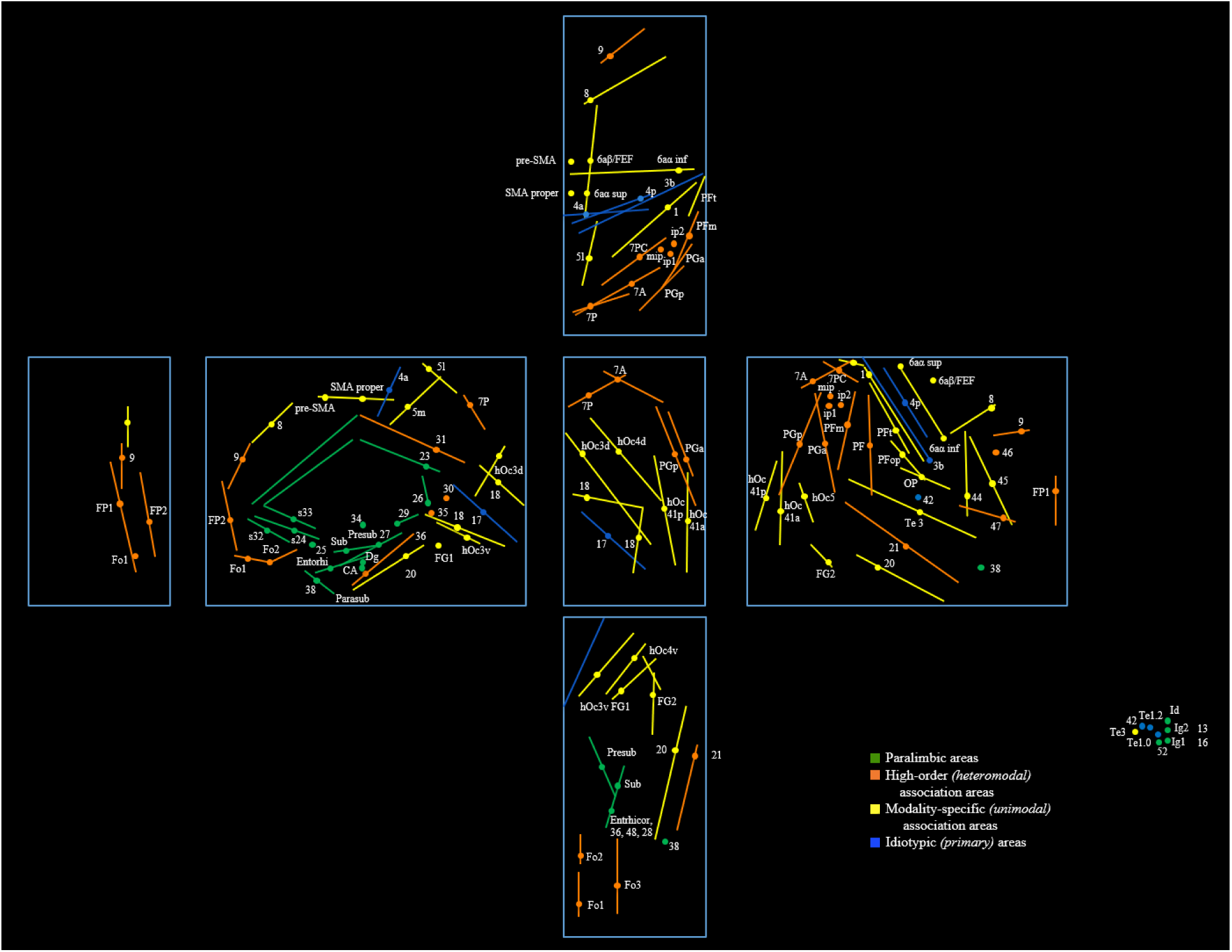
Mesulam^2,3^ hierarchies applicated to the Brodmann^40,41^/Zilles^45-47^-cytoarchitecture. The right hemisphere of a normalized^28,29^ brain, unfolded. Monofocal histological regions named following Brodmann & Zilles. FG3, FG4, 2, hip3, Te2.1, Te2.2, TPJ, TI, HATA, amygdala, ID2-3, OP5-7, p24a, p24b, pd24sv, pd24cv, pd24cd, pv24c, s24b, s24a, 25p, 25a, p32, CH1-4 anounced but not published. 8, 9, 38, 25, 34, 29, 26, 20, 35, 30, 23, 31, 24, 46, 47, 52 Brodmann-areas only. The points give the centers of gravity and the lines give the long axes of the areas volumes out of figure IV. The colors represent hierarchical hebbian layers, mono- or bisynaptic apart from each other.

**Figure VI.**
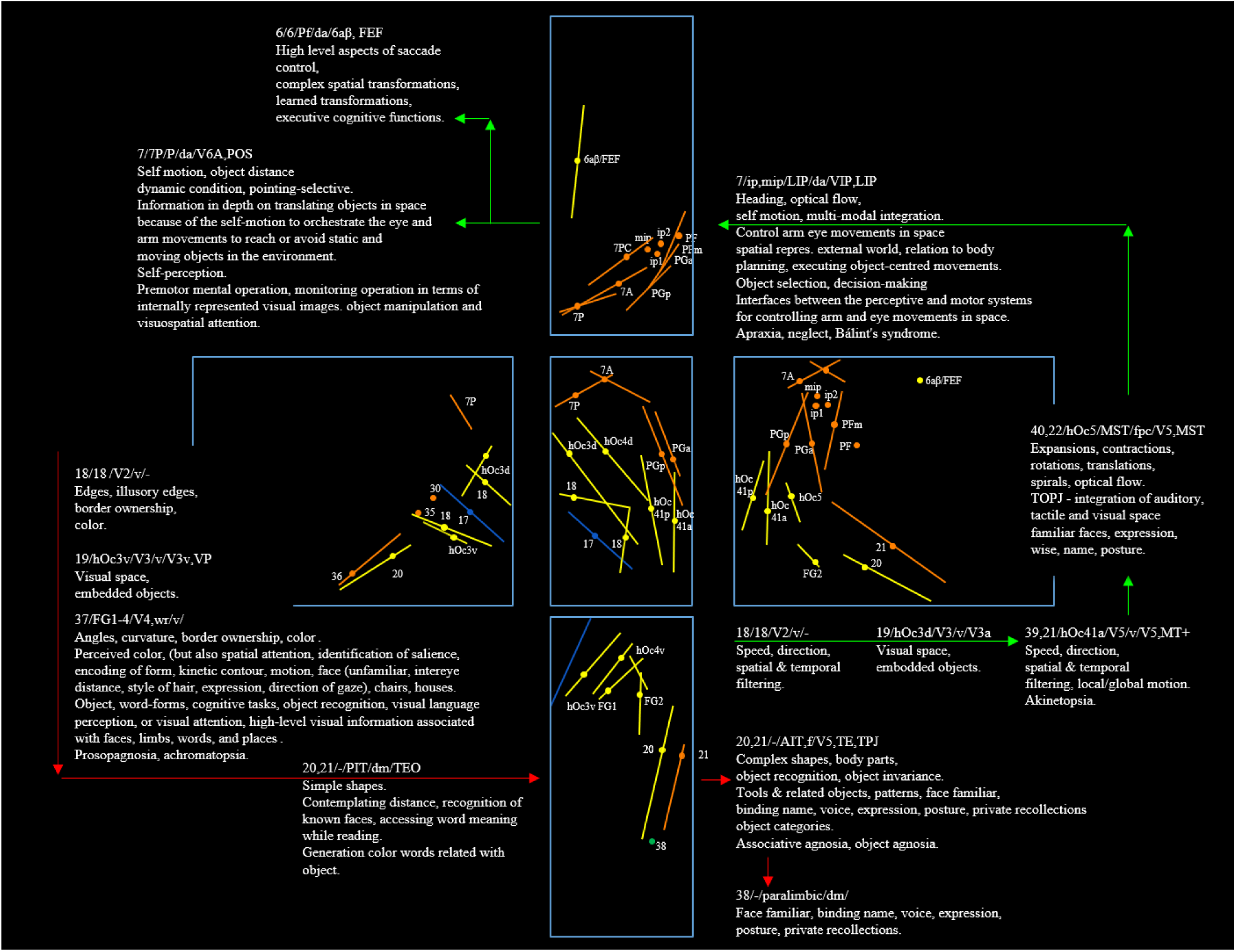
The architecture of the ventral and dorsal visual stream. Sculptured out of figure V. Paragraphs: top row - concordance between Brodmann^40,41^/Zilles^45-47^/Mesulam^2^/dfMRI^62,64-66,70,72,75,78-81^/other; following rows - presumed functions^2,56,57,59,60,62,61,58,91,97,104,108,105^, all using different number and localizations of nodes, as well as different numbers of hierarchical levels; bottom row - deficit-syndroms. Red - *The ventral stream … shows a considerable functional heterogeneity from early visual processing (posterior) to higher, domain-specific processing (anterior)*^*91*^, an ascending hierarchy with the sequence VI – V2 (V3) – V4 – inferior temporal cortex (IT), *starting with simpler features posteriorly (TEO/PIT) that increase in complexity as processing moves anteriorly (TE/AIT) to perform object recognition*^92^, asymmetrical, tending to the right hemisphere^93^: 17^94^ - V1, primary visual, retinotopic two-dimensional picturing of the external world; 18^94^ - gyrus lingualis, occipitalis inferior; hOc3v^95^ - lingual cortex; FG1/FG2^92,96^ - fusiform cortex, orientation selective cells concerned with form, and direction selective cells concerned with motion. *The fusiform gyrus hosts several “high-level” functional areas. It is involved in a core network subserving different cognitive tasks, that is, object recognition, visual language perception, or visual attention. It processes high-level visual information associated with faces, limbs, words, and places. FG2 is even a higher-order one*^96^; Green - The dorsal stream comprises the sequence V1 – V2 (V3) – middle temporal cortex, MT, V5, always retinotopic – medial superior temporal cortex, MST – and finally *intermediate object representations used by the decision-making circuitry further down the dorsal stream in lateral intraparietal sulcus, LIP*^97^; hOc3d^98^, cuneus, V3, V3A. There is a medio-temporal component consisting of hOc5^99,100^ - temporo-occipito-parietal TOP-junction, related to the deficit symptom of neglect^101^; and BA 21 - medial temporal gyrus, *it has been connected with processes as different as contemplating distance, recognition of known faces, and accessing word meaning while reading*^102^; *There is evidence to suggest that there is integration of both dorsal and ventral stream information into motion computation processes, giving rise to intermediate object representations, which facilitate object selection and decision-making mechanisms*^103^. *The areas within the intraparietal sulcus, IPS, in particular, serve as interfaces between the perceptive and motor systems for controlling arm and eye movements in space*^104^. This region is functionally and histologically complex in so as far as several association areas, default mode, executive control and dorsal attention^105,106^ overlap in here as well as the cytoarchitectonic areas 7A, 7PC, mip, Ip1, Ip2, PF and PFm, sulcus intraparietalis, *concerned with the integration of multimodal information for constructing a spatial representation of the external world in relation to the body or parts thereof, and planning and executing object-centered movements*^104^; 7P - precuneus, integration to a complex space out of proprioceptive and visual sources, *information on objects in depth which are translating in space, because of the self-motion, object distance in a dynamic condition (as that created by self-motion) to orchestrate the eye and arm movements necessary to reach or avoid static and moving objects in the environment*^107^, self-perception^108^. *Premotor area engaged in the mental operation, the precuneus aids monitoring the operation in terms of internally represented visual images. Interfaces between the perceptive and motor systems for controlling arm and eye movements in space. Visuomotor tasks comprising target selections for arm and eye movements, object manipulation and visuospatial attention, deficit syndrome of apraxia, neglect, Bálint’s syndrome*^109^.

**Figure VII.**
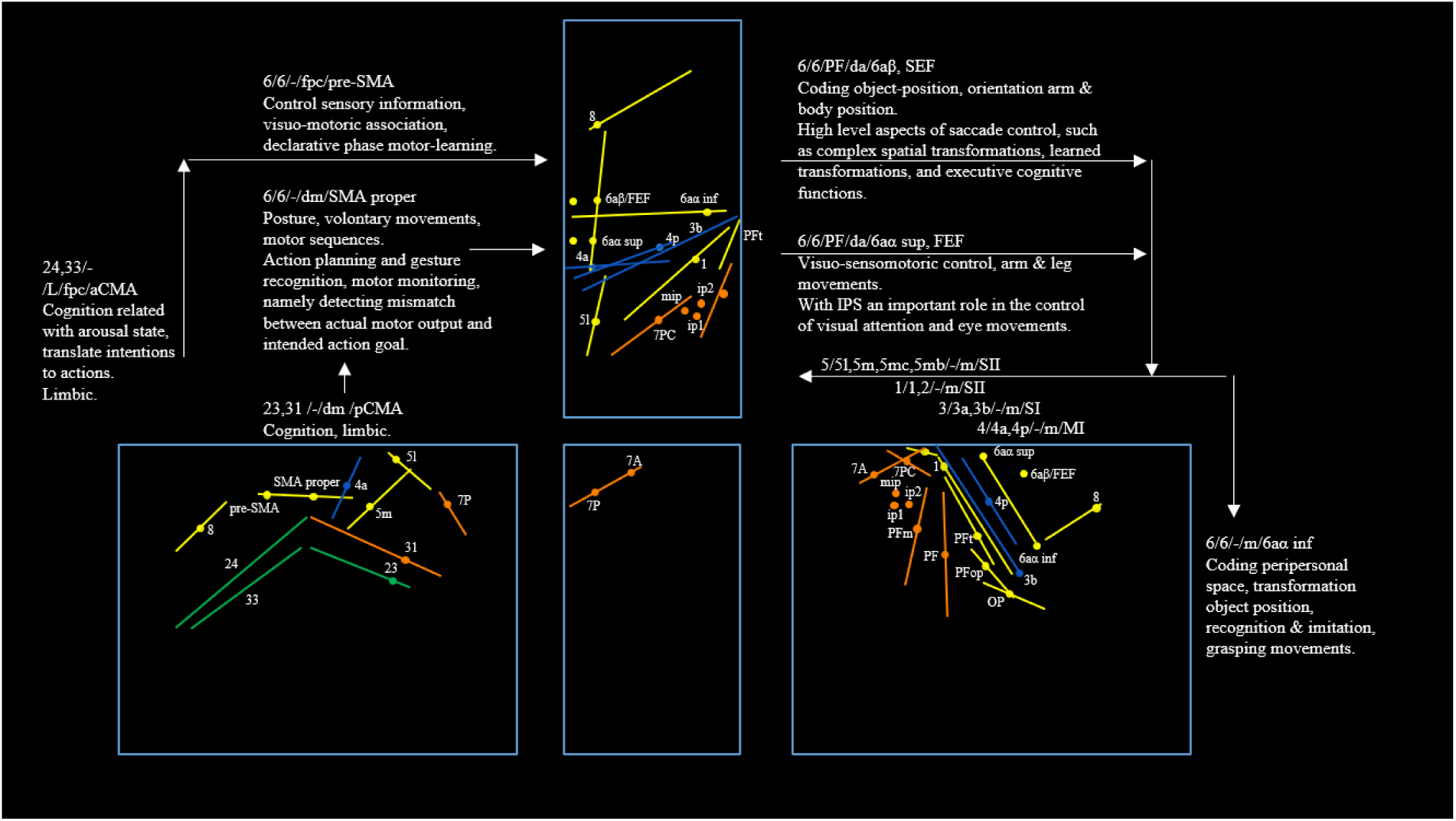
The sensorimotor architecture. Sculptured out of figure V. Paragraphs: top row - concordance between Brodmann^40,41^/Zilles^45-47^/Mesulam^2^/dfMRI^62,64-66,70,72,75,78-81^/other; following rows - presumed functions^12,60,104,110,118^. aCMA - cingulum anterior, *its proposed role in cognition and its relationship with the arousal/drive state of the organism placing it in a unique position to translate intentions to actions*^110^, and being dedicated to the limbic portion of the cortex; 6^111^ - pre-SMA and SMA proper^112,113^, intention to act, generation of readiness potential^114,115,116,12^, *action planning and gesture recognition, the operation of the motor monitoring, namely detecting the mismatch between the actual motor output and intended action goal*^117^. *SMA proper and pre-SMA are active prior to volitional movement or action, as well as the cingulate motor area (CMA). This is now known as ‘anterior mid-cingulate cortex (aMCC)’. It has been shown by integrating simultaneously acquired EEG and fMRI that SMA and aMCC have strong reciprocal connections that act to sustain each other’s activity, and that this interaction is mediated during movement preparation according to the Bereitschaftspotential amplitude*^112^; 6aβ - the supplementary eye field (SEF), which has a special role in high level aspects of saccade control, such as complex spatial transformations, learned transformations, and executive cognitive functions^118,119^; 6aα superior - visuo- somatomotor area hosting the frontal eye field (FEF)^120^, *which constitutes together with the intraparietal sulcus (IPS)* … *an important role in the control of visual attention and eye movements*^*104*^, the coding of object positions^59^; 4p^121^ - central sulcus, primary area, *involved in kinematic and dynamic parameters of voluntary movements*^121^; 3a/3b,1^122^ - primary area; 2^123^ - the projection emphasising size and shape, secondary unimodal; 5^124^ - secondary, unimodal.

**Figure VIII.**
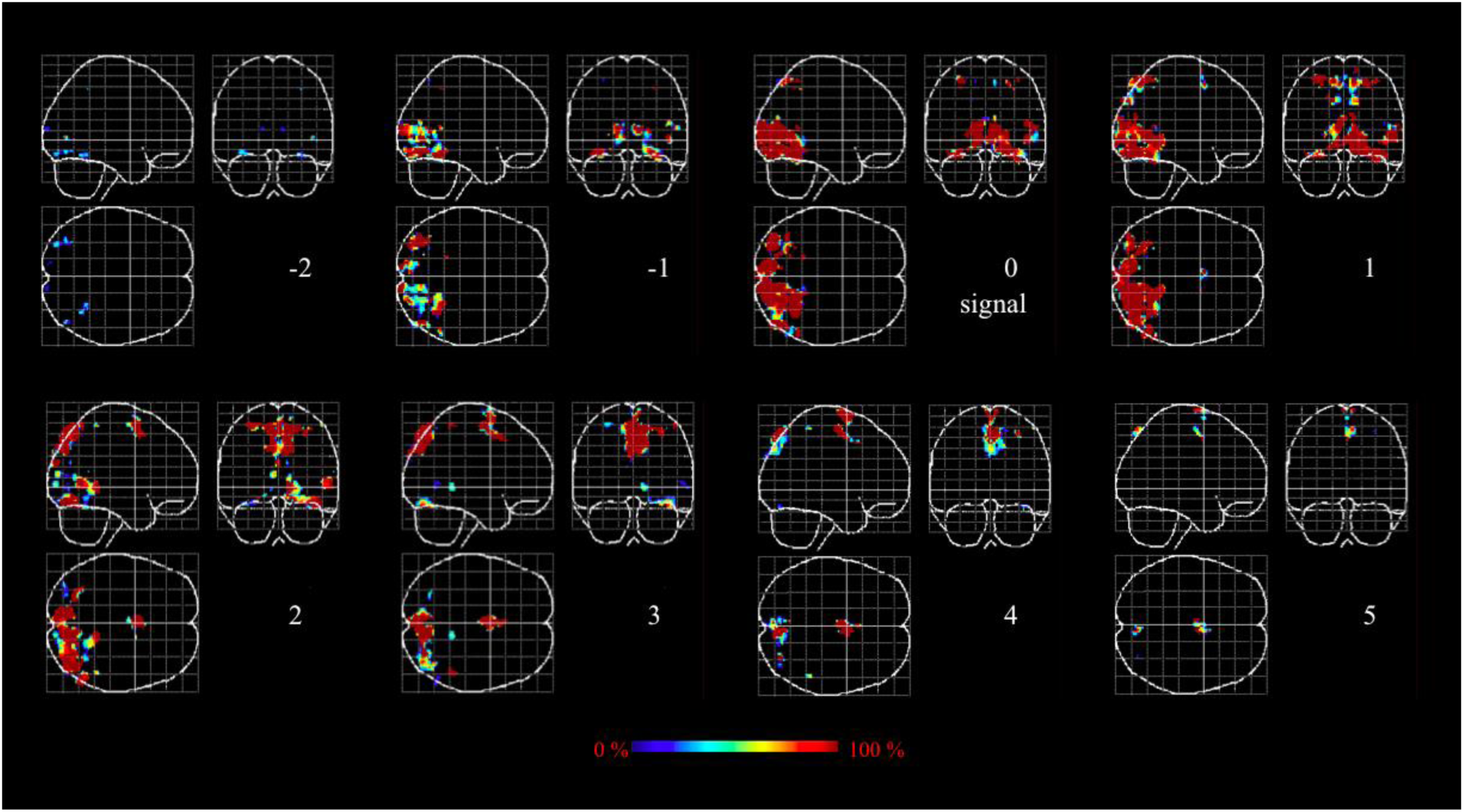
Perception-related time series. Time steps in second distances (2 seconds ante until 5 seconds post perception, white numbers). Perception at time step three. Activity strength following the colour bar (see supplementary movie II, part i).

**Figure IX.**
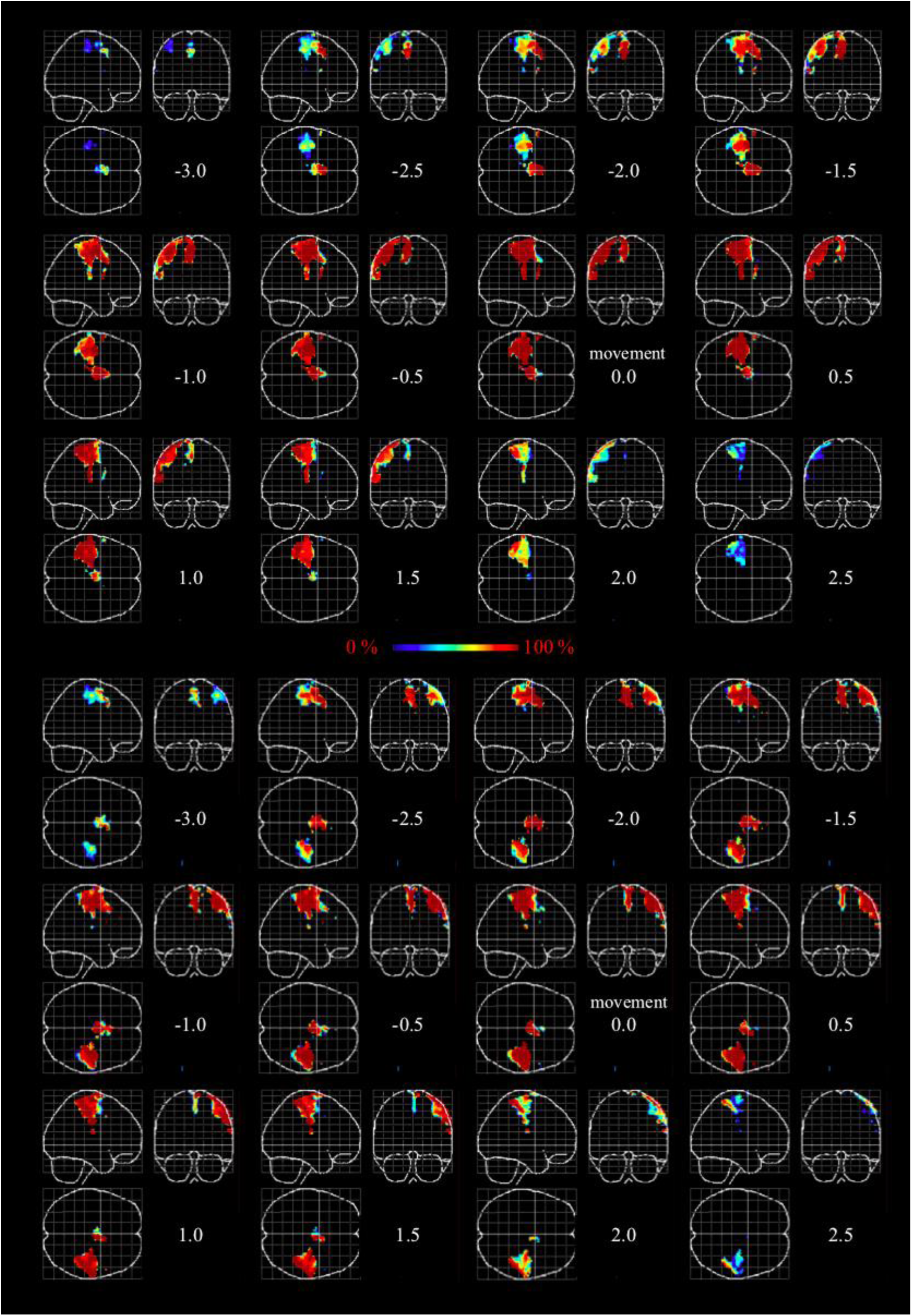
Reaction-related time series. Right hand - above; left hand - below. Time steps in half second distances (3 seconds ante until 2.5 seconds post reaction, white numbers). Reaction at time step eight. Activity strength following the colour bar (see supplementary movie II, part ii).

**Figure X.**
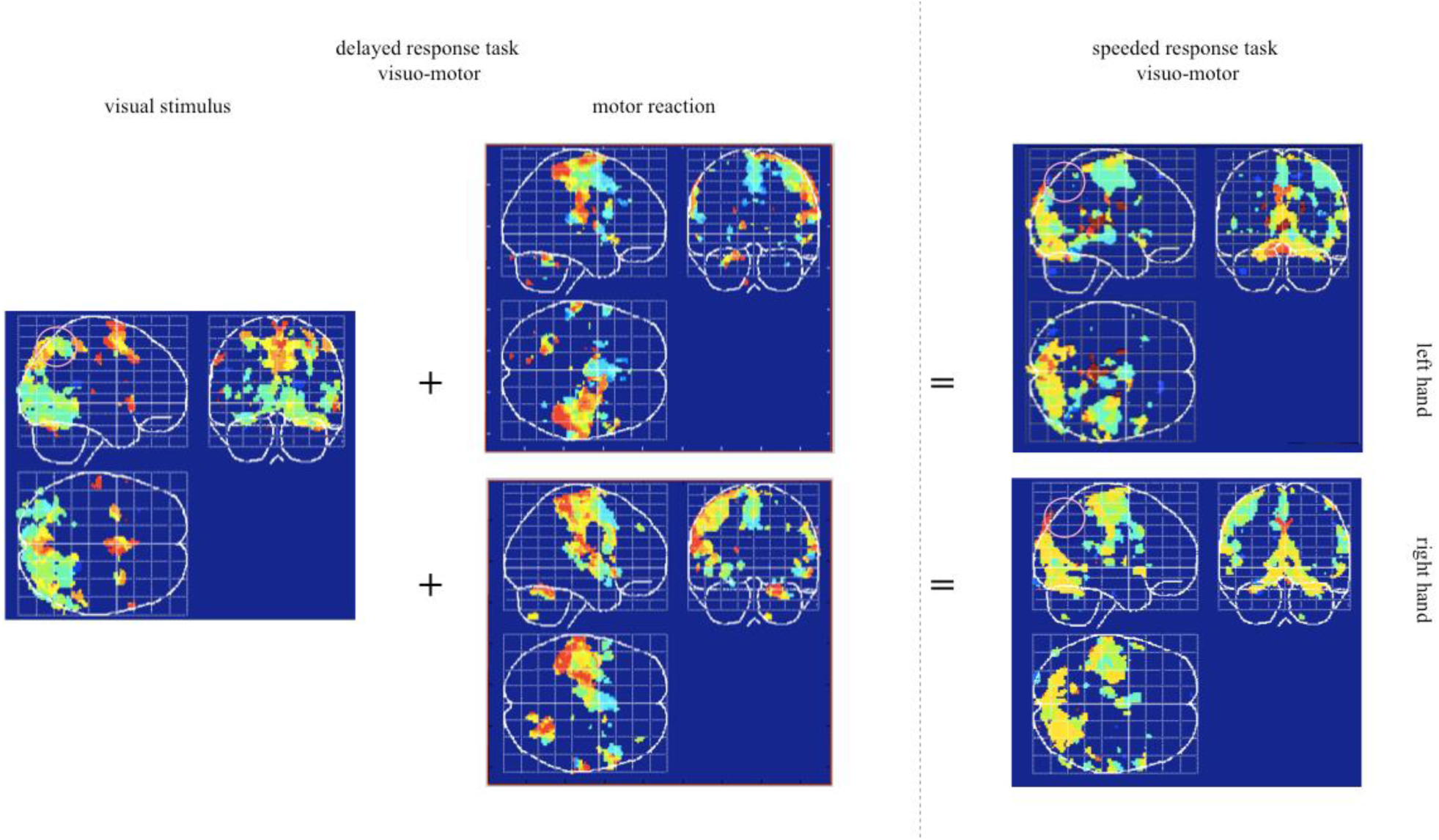
Contrast of delayed and speeded responses, phase images. The unique hierarchical position of the complex made of the parietal Brodmann area 7, a core area of the dorsal attention network^62,70,72,75,78-81^ becomes clear in contrast imaging of speeded and delayed-response tasks. The speeded responses (right) activate all of the same regions as the sum of the delayed responses (the visual phase image on the left, next to it both motor phase images) – with the exception of the high-association superior parietal regions (precuneus and parietalis superior, BA 7, pink rings). The high-association areas are apparently used only when they are involved in long-lasting feedforward-feedback-loops or, in short, in the case of accelerated responses, there is no time left for daydreaming.

Movie I shows the Mesulam model unfolded over time.

## SUPPLEMENTARY

**Movie I.**
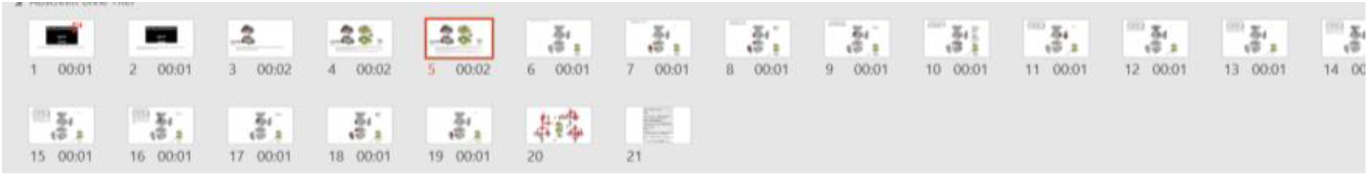
Traveling cortical Netwave, hypothesis^125^. https://youtu.be/cQVBLAThJqk Jubrain^45-47^ regions aligned over time and neighbourhood from the visual perception to the voluntary motor reaction, and back over the autoperception and heteromodal association, assigned to the cortical hierarchies (coloured box bottom, right) of Mesulam^2^, unfolded right cortical hemisphere.

**Movie II.**
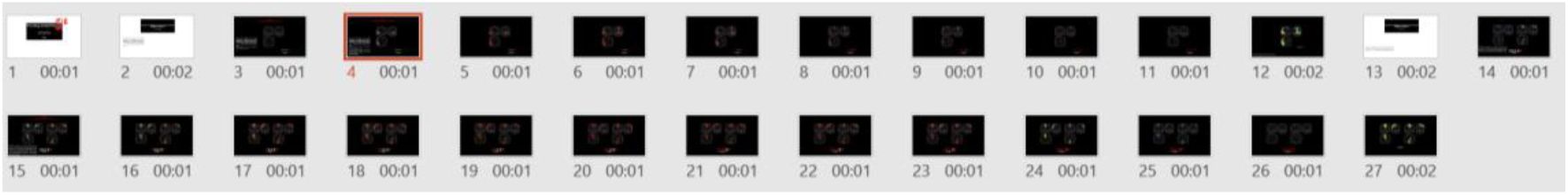
Traveling cortical Netwave, results^126^. https://youtu.be/efEaO79gBkM Part i - The time series for the visual perception, 2 seconds ante until 5 seconds post perception, perception at t = 0 s; last picture the resulting phase image. Part ii - The right and left hand motor reaction, 3.5 seconds ante until 2 seconds post reaction, movement at t = 0 s; last picture the resulting phase image. The red-white boxes at the bottom represent the time axis in seconds. Activity strength (scaled t-values of hrf-correlation) following the colour bar at the bottom, but in frame 12 and 27, the resulting phase images, where colour codes time.

**Movie III.**
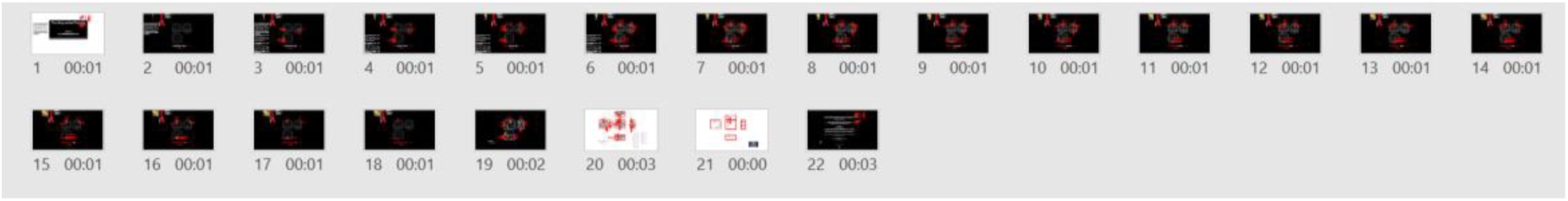
Traveling cortical Netwave, interpretation^127^. https://youtu.be/fvLWPrw2Ro0 The fMRI time series for the visual perception and the following the left hand motor act, fusioned out of sup. movie II, using the medial supplementary motor area (resulting time axis 2 seconds ante until 7.5 seconds post perception, 7 seconds ante until 2.5 seconds post reaction), The red-white boxes at the bottom represent the time axis, scaled in seconds above with respect to the visual signal and below with respect to the motor reaction. Activity strength (scaled t-values of hrf-correlation) following the colour bar at the bottom. The anatomical regions labelled following Jubrain^45-47,91,94-96,98,99,100,109^. The evolution of different standing netwaves, the resulting travelling netwave and winning coalitions^5^ following AAL^32^ assigned to the hierarchies of Mesulam^2,3^ in the insert above left (see sup. tab. III). Frame 20 gives the surface rendered phase image, right hemisphere, as well as the extreme values in space and time (left table) and the resulting vector-matrix (right table). Colour codes time, with respect to the signal (upper timescale) and to the reaction (lower timescale). Labelling JuBrain. Temporo-spatial cortical vectors red. Their position in space (x, y, z, Talairach-space (blue boxes) in mm) and time (t in s) as shown in the left table. Their length in time (dt in s) and space (ds in mm) in the right table.

## Notes

https://youtu.be/cQVBLAThJqk

https://youtu.be/efEaO79gBkM

https://youtu.be/fvLWPrw2Ro0

### Competing Interest Statement

The authors have declared no competing interest.

## REFERENCES

1. Hebb D. The Organization of Behavior. Wiley New York (1949).

2. Mesulam MM. From sensation to cognition. Brain 121, 1013–1052 (1998).

3. Mesulam MM. Principles of Behavioral and Cognitive Neurology. Oxford University Press (2000).

4. Deco G. Tononi G. Boly M. Kringelbach ML. Rethinking segregation and integration: contributions of whole-brain modelling. Nat Rev Neurosci. 16, 430–439 (2015).

5. Crick F. Koch C. A framework of consciousness. Nat Neurosci. 6, 119–126 (2003).

6. Deco G. Jirsa VK. Robinson PA. Breakspear M. Friston K. The dynamic brain: from spiking neurons to neural masses and cortical fields. PLoS Comput Biol. 4, (2008).

7. Nieuwenhuys R. Donkelaar HJ. Nicholson C. The central nervous system of vertebrates, Volume 1. Springer, 2011–2012 (1998).

8. Toro R. Perron M. Pike B. Richer L. Veillette S. Pausova Z. Paus T. Brain Size and Folding of the Human Cerebral Cortex. Cerebral Cortex 18, 2352–2357 (2008).

9. Haynes JD. Rees G. Decoding mental states from brain activity in humans. Nature Reviews Neuroscience 7, 523–534 (2006).

10. He BJ. Raichle ME. The fMRI signal, slow cortical potential and consciousness. Trends Cogn Sci.13, 302–309 (2009).

11. Birbaumer N. Elbert T. Canavan AG. Rockstroh B. Slow potentials of the cerebral cortex and behavior. Physiol Rev. 70, 1–41 (1990).

12. Montemurro MA. Rasch MJ. Murayama Y. Logothetis NK. Panzeri S. Phase-of-firing coding of natural visual stimuli in primary visual cortex. Curr. Biol 18, 375–380 (2008).

13. Lakatos P. Karmos G. Mehta AD. Ulbert I. Schroeder CE. Entrainment of neuronal oscillations as a mechanism of attentional selection. Science 320, 110–113 (2008).

14. Muller L. Chavane F. Reynolds J. Sejnowski TJ. Cortical travelling waves: mechanisms and computational principle. Nat Rev Neurosci.19, 255–268 (2018).

15. Calhoun V. Liégeois R. Time-varying Connectivity in Resting-state fMRI: Methods, interpretations and clinical use. OHBM (2019).

16. Lurie D. Controversies and Open Questions in the Study of “Resting State” Time-Varying Functional Connectivity. OHBM (2019).

17. Breakspear M. Modelling the Multiscale Nature of Dynamic Functional Connectivity. OHBM (2019).

18. Breakspear M. Terry JR. Friston KJ. Modulation of excitatory synaptic coupling facilitates synchronization and complex dynamics in a biophysical model of neuronal dynamics. Network 14, 703–732 (2003).

19. Erb M. Aersten A. Dynamics of Activity in Biology-Oriented Neural Network Models: Stability at Low Firing Rates. In: Information Processing in the Cortex. Experiment and Theory. Edited by Braitenberg V. & Aertsen A. Springer (1992).

20. Omidvarnia A. Pedersen M. Walz JM. Vaughan DN. Abbott DF. Jackson GD. Dynamic regional phase synchrony (DRePS): An Instantaneous Measure of Local fMRI Connectivity Within Spatially Clustered Brain Areas. Hum Brain Mapp. 37, 1970–1985 (2016).

21. Friston KJ. Ashburner JT. Kiebel SJ. Nichols TE. Penny WD. Statistical Parametric Mapping. Academic Press (2008).

22. Fornito A. Zalesky Z. Bullmore E. Fundamentals of Brain Network Analysis. Academic Press (2016).

23. Fernández-Corazza M. Turovets S. Luu P. Anderson E. Tucker D. Transcranial Electrical Neuromodulation Based on the Reciprocity Principle. Front. Psychiatry 27 May (2016).

24. Huelsmann E. Erb M. Grodd W. From will to action: sequential cerebellar contributions to voluntary movement. Neuroimage 20, 1485–1492 (2003).

25. Nolte J. (editor). The Human Brain: An Introduction to its functional Anatomy. Elsevier, 510–517 (2008).

26. Bordier C. Macaluso E. Time-resolved detection of stimulus/task-related networks, via clustering of transient intersubject synchronization. Hum Brain Mapp. 36, 3404–3425 (2015).

27. Cohen MX. Analysing Neural Time Series Data. The MIT Press (2014).

28. Talairach J. David M. Tournoux P. Atlas d’anatomie stéréotaxique. Masson Paris (1957).

29. Mazziotta JC. Toga AW. Evans AP. Fox A. Lancaster J. A Probablistic Atlas of the Human Brain: Theory and Rationale for Its Development. Neuroimage 2, 89–101 (1995).

30. Ecker A. Die Hirnwindungen des Menschen: nach eigenen Untersuchungen; insbesondere über die Entwicklung derselben beim Fötus und mit Rücksicht auf das Bedürfnis der Ärzte. Vieweg Braunschweig (1869).

31. Smith EG. A new topographical survey of the human cerebral cortex, being an account of the distribution of the anatomically distinct cortical areas and their relationship to the cerebral sulci. J. Anat. 41, 237–254 (1907).

32. Tzourio-Mazoyer N. Landeau B. Papathanassiou D. Crivello F. Etard O. Delcroix N. Mazoyer B. Joliot M. Automated anatomical labeling of activations in SPM using a macroscopic anatomical parcellation of the MNI MRI single-subject brain. Neuroimage 15, 273–289 (2002).

33. Makris, N. Goldstein JM. Kennedy D. Hodge SM. Caviness VS. Faraone SV. Tsuang MT. Seidman LJ. Decreased volume of left and total anterior insular lobule in schizophrenia. Schizophr. Res. 83, 155–171 (2006).

34. Desikan RS. Ségonne F. Fischl B. Quinn BT. Dickerson BC. Blacker D. Buckner RL. Dale AM. Maguire RP. Hyman BT. Albert MS. Killiany RJ. An automated labeling system for subdividing the human cerebral cortex on MRI scans into gyral based regions of interest. Neuroimage 31, 968–980 (2006).

35. Gousias IS. Rueckert D. Heckemann RA. Dyet LE. Boardman JP. Edwards AD. Hammers A. Automatic segmentation of brain MRIs of 2-year-olds into 83 regions of interest. NeuroImage 40, 672–684 (2008).

36. Destrieux C. Fischl B. Dale A. Halgren E. Automatic parcellation of human cortical gyri and sulci using standard anatomical nomenclature. Neuroimage 53, 1–15 (2010).

37. Klein A. Tourville J. 101 labeled brain images and a consistent human cortical labeling protocol. Frontiers in Neuroscience 6, 171 (2012).

38. Auzias G. Coulon O. Brovelli A. MarsAtlas: A cortical parcellation atlas for functional mapping. Human Brain Mapping 37, 1573–1592 (2016).

39. Campbell AW. Histological Studies on the Localisation of Cerebral Function. Cambridge Univ. Press, Cambridge, UK (1905).

40. Brodmann K. Vergleichende Lokalisationslehre der Großhirnrinde in ihren Prinzipien dargestellt auf Grund ihres Zellenbaues. Barth, Leipzig (1909).

41. Lacadie CM. Fulbright RK. Arora J. Constable RT. Papademetris X. Brodmann Areas defined in MNI space using a new Tracing Tool in BioImage Suite. Human Brain Mapping, (2008).

42. von Economo C. Koskinas GN. Die Cytoarchitektonik der Hirnrinde des erwachsenen Menschen. Springer Verlag Wien (1925).

43. Sarkisov SA. Filimonoff IN. Preobrashenskaya NS. Cytoarchitecture of the human cortex cerebri. Medgiz Moscow (1949).

44. Bailey P. von Bonin G. The Isocortex of Man. Univ. Illinois Press, Urbana (1951).

45. Amunts K. Zilles K. Architectonic Mapping of the Human Brain beyond Brodmann. Neuron 16, 1086–1107 (2015).

46. Eickhoff SB. Stephan KE. Mohlberg H. Grefkes C. Fink GR. Amunts K. Zilles K. A new SPM toolbox for combining probabilistic cytoarchitectonic maps and functional imaging data. Neuroimage 25, 1325–1335 (2005).

47. Mohlberg H. Eickhoff SB. Schleicher A. Zilles K. Amunts K. A new processing pipeline and release of cytoarchitectonic probabilistic maps –JuBrain. OHBM Peking, China (2012).

48. Vogt C. Vogt O. Allgemeine Ergebnisse unserer Hirnforschung. J Psychol Neural. 25, 277–462 (1919).

49. Flechsig PE. Anatomie des menschlichen Gehirns und Rückenmarks auf myelogenetischer Grundlage, Volume 1. Anatomie des menschlichen Gehirns und Rückenmarks auf myelogenetischer Grundlage. G. Thieme (1920).

50. Geschwind N. Disconnexion syndromes in animals and man. I. Brain 88, 237–294 (1965).

51. Catani M. Ffytch DH. The rises and falls of disconnection syndromes. Brain 128, 2224–2239 (2005).

52. Van Essen DC. Ugurbil K. Auerbach E. Barch D. Behrens TE. Bucholz R. Chang A. Chen L. Corbetta M. Curtiss SW. Della Penna S. Feinberg D. Glasser MF. Harel N. Heath AC. Larson-Prior L. Marcus D. Michalareas G. Moeller S. Oostenveld R. Petersen SE. Prior F. Schlaggar BL. Smith SM. Snyder AZ Xu J. Yacoub E. The Human Connectome Project: a data acquisition perspective. Neuroimage 62, 2222–2231 (2012).

53. Gall FJ. Philosophisch-medizinische Untersuchungen über Natur und Kunst im kranken und gesunden Zustand des Menschen. Gräffer, Wien (1791).

54. Zilles K. Rehkaemper G. Funktionelle Neuroanatomie. Springer (1993).

55. Van Essen DC. Drury HA. Joshi S. Miller MI. Functional and structural mapping of human cerebral cortex: solutions are in the surfaces. Adv Neurol. 84, 23–34 (2000).

56. Van Essen DC. Anderson CH. Felleman DJ. Information processing in the primate visual system: an integrated systems perspective. Science 255, 419–423 (1992).

57. Ungerleider LG. Functional brain imaging studies of cortical mechanisms for memory. Science 270, 769–775 (1995).

58. Glerean E. Salmi J. Lahnakoski JM. Iiro P. Jääskeläinen IP. Mikko Sams M. Functional Magnetic Resonance Imaging Phase Synchronization as a Measure of Dynamic Functional Connectivity. Brain Connect. 2, 91–101 (2012).

59. Naser K. Ricordel V. Le Callet P. in Benois-Pineau J. Le Callet P. Visual content indexing and retrieval with psycho-visual models. Springer, 20 (2017).

60. Grandinetti L. Lippert T. Petkov N. Brain-inspired computing. *Springer,* 53 (2013).

61. Pitzalis S. Fattori P. Galletti C. The human cortical areas V6 and V6A. Vis Neurosci., 32 (2015).

62. Yeo BT. Krienen FM. Sepulcre J. Sabuncu MR. Lashkari D. Hollinshead M. Roffman JL. Smoller JW. Zollei L. Polimeni JR. Fischl B. Liu H. Buckner RL. The organization of the human cerebral cortex estimated by intrinsic functional connectivity. J Neurophysiol. 106, 1125–1165 (2011).

63. Zilles K. Schlaug G. Matelli M. Luppino G. Schleicher A. Qü M. Dabringhaus A. Seitz R. Roland PE. Mapping of human and macaque sensorimotor areas by integrating architectonic, transmitter receptor, MRI and PET data. J Anat. 187, 515–537 (1995).

64. Smith SM. Fox PT. Miller KL. Glahn DC. Fox PM. Mackay CE. Filippini N. Watkins KE. Toro R. Laird AR. Beckmann CF. Correspondence of the brain’s functional architecture during activation and rest. Proc Natl Acad Sci USA. 106, 13040–13045 (2009).

65. Andrews-Hanna JR. Smallwood J. Spreng RN. The default network and self-generated thought: component processes, dynamic control, and clinical relevance. Ann N Y Acad Sci. 1316, 29–52 (2014).

66. Andrews-Hanna JR. Reidler JS. Sepulcre J. Poulin R. Buckner RL. Functional-anatomic fractionation of the brain’s default network. Neuron 65, 550–562 (2010).

67. Bellec P. Rosa-Neto P. Lyttelton OC. Benali H. Evans AC. Multi-level bootstrap analysis of stable clusters in restingstate fMRI. NeuroImage 51, 1126–1139 (2010).

68. Power JD. Cohen AL. Nelson SM. Wig GS. Barnes KA. Church JA. Vogel AC. Laumann TO. Miezin FM. Schlaggar BL. Petersen SE. Functional network organization of the human brain. Neuron 72, 665–678 (2011).

69. Buckner RL. Krienen FM. Castellanos A. Diaz JC. Yeo BT. The organization of the human cerebellum estimated by intrinsic functional connectivity. J. Neurophysiol. 106, 2322–2345 (2011).

70. Buckner RL. Krienen FM. Yeo BTT. Opportunities and limitations of intrinsic functional connectivity MRI. Nature Neuroscience. 16, 832–837 (2013).

71. Choi EY. Yeo BT. Buckner RL. The organization of the human striatum estimated by intrinsic functional connectivity. J Neurophysiol. 108, 2242–2263 (2012).

72. Choi EY. Drayna GK. Badre D. Evidence for a Functional Hierarchy of Association Networks. J Cogn Neurosci. 30, 722–736 (2018).

73. Craddock RC. James GA. Holtzheimer PE. Hu XP. Mayberg HS. A whole brain fMRI atlas generated via spatially constrained spectral clustering. Human Brain Mapping 33, 1914–1928 (2012).

74. Shen X. Tokoglu F. Papademetris X. Constable, RT. Groupewise whole-brain parcellation from resting-state fMRI data for network node identification. NeuroImage 82, 403–415 (2013).

75. Spreng RN. Sepulcre J. Turner GR. Stevens WD, Schacter DL. Intrinsic architecture underlying the relations among the default, dorsal attention, and frontoparietal control networks of the human brain. J Cogn Neurosci. 25, 74–86 (2013).

76. Wang D. Buckner RL. Fox MD. Holt DJ. Holmes AJ. Stoecklein S. Langs G. Pan R. Qian T. Li K. Baker JT. Stufflebeam SM. Wang K. Wang X. Hong B. Liu H. Parcellating cortical functional networks in individuals. Nat Neurosci. 18, 1853–1860 (2015).

77. Joliot M. Jobard G. Naveau M. Delcroix N. Petit L. Zago L. Crivello F. Mellet E. Mazoyer B. Tzourio-Mazoyer N. AICHA: An atlas of intrinsic connectivity of homotopic areas. Journal of Neuroscience Methods 254, 46–59 (2015).

78. Zhang D. Liang B. Wu X. Wang Z. Xu P. Chang S. Liu B. Liu M. Huang R. Directionality of large-scale resting-state brain networks during eyes open and eyes closed conditions. Front Hum Neurosci. 9, 81 (2015).

79. Fan L. Li H. Zhuo J. Zhang Y. Wang J. Chen L. Yang Z. Chu C. Xie S. Laird AR. Fox PT. Eickhoff SB. Yu C. Jiang T. The Human Brainnetome Atlas: A New Brain Atlas Based on Connectional Architecture. Cerebral Cortex 26, 3508–3526 (2016).

80. Gordon EM. Laumann TO. Adeyemo B. Huckins JF. Kelley WM. Petersen SE. Generation and Evaluation of a Cortical Area Parcellation from Resting-State Correlations. Cerebral Cortex 26, 288–303 (2016).

81. Braga RM. Buckner RL. Parallel interdigitated distributed networks within the individual estimated by intrinsic functional connectivity. Neuron 95, 457–472 (2017).

82. Schaefer A. Kong R. Gordon EM, Laumann TO. Zuo XN. Holmes AJ. Eickhoff SB. Yeo BTT. Local-global parcellation of the human cerebral cortex from intrinsic functional connectivity MRI. Cerebral Cortex 28, 3095–3114 (2017).

83. Kong R. Li J. Sun N. Sabuncu MR. Liu H. Schaefer A. Orban C. Zuo XN. Holmes AJ. Eickhoff S. Yeo B.T. Spatial topography of individual-specific cortical networks predicts human cognition, personality and emotion. Cerebral Cortex https://doi.org/10.1093/cercor/bhy123 (2018).

84. Huth AG. de Heer WA. Griffiths TL. Theunissen FE. Gallant JL. Natural speech reveals the semantic maps that tile human cerebral cortex. Nature 532, 453–458 (2016).

85. Huth AG. Griffiths TL. Theunissen FE. Gallant JL. PrAGMATiC: a probabilistic and generative model of areas tiling the cortex. arXiv (2015).

86. Fan Y. Zeng LL. Shen H. Qin J. Li F. Hu D. Lifespan Development of the Human Brain Revealed by Large-Scale Network Eigen-Entropy. Entropy 19, 471 (2017).

87. Glasser MF. Coalson TS. Robinson EC. Hacker CD. Harwell J. Yacoub E. Ugurbil K. Andersson J. Beckmann CF. Jenkinson M. Smith SM. Van Essen DC. A multi-modal parcellation of human cerebral cortex. Nature 536, 171–178 (2016).

88. Varikuti DP. Genon S. Sotiras A. Schwender H. Hoffstaedter F. Patil KR. Jockwitz C. Caspers S. Moebus S. Amunts K. Davatzikos C. Eickhoff SB. Evaluation of non-negative matrix factorization of grey matter in age prediction. NeuroImage 173, 394–410 (2018).

89. Eickhoff SB. Yeo BTT. Genon S. Imaging-based parcellations of the human brain. Nat Rev Neurosci. 19, 672–686 (2018).

90. Logothetis NK. The neural basis of the blood-oxygen-level-dependent functional magnetic resonance imaging signal. Philos. Trans. R. Soc. Lond. B Biol. Sci. 357, 1003–1037 (2002).

91. Caspers J. Zilles K. Amunts K. Laird AR. Fox PT. Eickhoff SB. Functional characterization and differential coactivation patterns of two cytoarchitectonic visual areas on the human posterior fusiform gyrus. Hum Brain Mapp. 6, 2754–2767 (2014).

92. Cowey A. Weiskrantz L. A comparison of the effects of inferotemporal and striate cortex lesions on the visual behaviour of rhesus monkeys. J Exp Psychol. 3, 246–253 (1967).

93. Karnath HO. Insights into the functions of the superior temporal cortex. Nat Rev Neurosci. 2, 568–575 (2001).

94. Amunts K. Malikovic A. Mohlberg H. Schormann T. Zilles K. Brodmann’s areas 17 and 18 brought into stereotaxic space-where and how variable? Neuroimage 11, 66–84 (2000).

95. Rottschy C. Eickhoff SB. Schleicher A. Mohlberg H. Kujovic M. Zilles K. Amunts K. Ventral visual cortex in humans: cytoarchitectonic mapping of two extrastriate areas. Human Brain Mapp. 28, 1045–1059 (2007).

96. Lorenz S. Weiner KS. Caspers J. Mohlberg H. Schleicher A. Bludau S. Eickhoff SB. Grill-Spector K. Zilles K. Amunts K. Two New Cytoarchitectonic Areas on the Human Mid-Fusiform Gyrus. Cerebral Cortex. 27, 373–385 (2017).

97. Van Polanena V. Davarea M. Interactions between dorsal and ventral streams for controlling skilled grasp. Neuropsychologia 79, 186–191 (2015).

98. Kujovic M. Zilles K. Malikovic A. Schleicher A. Mohlberg H. Rottschy C. Eickhoff SB. Amunts K. Cytoarchitectonic mapping of the human dorsal extrastriate cortex. Brain Struct Funct. 218, 157–172 (2013).

99. Malikovic A. Amunts K. Schleicher A, Mohlberg H. Eickhoff SB. Wilms M. Palomero-Gallagher N. Armstrong E. Zilles K. Cytoarchitectonic analysis of the human extrastriate cortex in the region of V5/MT+: a probabilistic, stereotaxic map of area hOc5. Cereb Cortex 17, 562–574 (2007).

100b. Malikovic A. Amunts K. Schleicher A. Mohlberg H. Kujovic M. Palomero-Gallagher N. Eickhoff SB. Zilles K. Cytoarchitecture of the human lateral occipital cortex: mapping of two extrastriate areas hOc4la and hOc4lp. Brain Structure and Function, 221, 1877–1897 (2016).

100. Caspers S. Eickhoff SB. Geyer S. Scheperjans F. Mohlberg H. Zilles K. Amunts K. The human inferior parietal lobule in stereotaxic space. Brain Struct Funct. 212, 481–495 (2008).

101. Vallar G. Spatial hemineglect in humans. Trends Cogn Sci. 2, 87–95 (1998).

102. Acheson DJ. Hagoort P. Stimulating the brain’s language network: syntactic ambiguity resolution after TMS to the inferior frontal gyrus and middle temporal gyrus. J Cogn Neurosci., 1664–1677 (2013).

103. Perry CJ. Fallah M. Feature integration and object representations along the dorsal stream visual hierarchy. Front Comput Neurosci. 8, 84 (2014).

104. Grefkes C. Fink GR. The functional organization of the intraparietal sulcus in humans and monkeys. J Anat. 207, 3–17 (2005).

105. Oshio R, Tanaka S, Sadato N, Sokabe M, Hanakawa T, Honda M. Differential effect of double-pulse TMS applied to dorsal premotor cortex and precuneus during internal operation of visuospatial information. Neuroimage 49, 1108–1115 (2010).

106. Kjaer TW. Nowak M. Lou HC. Reflective self-awareness and conscious states: PET evidence for a common midline parietofrontal core. Neuroimage 17, 1080–1086 (2002).

107. Pitzalis S. Fattori P. Galletti C. The human cortical areas V6 and V6A. Vis Neurosci. 32 (2015).

108. Birbaumer N. Sanyung K. Dein Gehirn weiss mehr als Du denkst. Ullstein, 230–233 (2016).

109. Choi HJ. Zilles K. Mohlberg H. Schleicher A. Fink GE. Armstrong E. Amunts K. Cytoarchitectonic identification and probabilistic mapping of two distinct areas within the anterior ventral bank of the human intraparietal sulcus. J Comp Neurol. 495, 53–69 (2006).

110. Paus T. Primate anterior cingulate cortex: Where motor control, drive and cognition interface. Nat Rev Neurosci. 2, 417–424 (2001).

111. Geyer S. Matelli M. Luppino G. Zilles K. Functional neuroanatomy of the primate isocortical motor system Anat Embryol. 202, 443–474 (2000).

112. Nguyen VT. Breakspear M. Cunnington R. Reciprocal interactions of the SMA and cingulate cortex sustain pre-movement activity for voluntary actions. J Neurosci. 34, 16397–16407 (2014).

113. Picard N. PL. Motor areas of the medial wall: a review of their location and functional activation. Cerebral Cortex 6, 342–53 (1996).

114. Libet B. Gleason CA. Wright EW. Pearl DK. Time of conscious intention to act in relation to onset of cerebral activity (readiness-potential). The unconscious initiation of a freely voluntary act. Brain 106, 623–842 (1983).

115. Kornhuber HH. Deecke L. Changes in the brain potential in voluntary movements and passive movements in mean readiness potential and reafferent potentials. Pflugers Arch Gesamte Physiologie Menschen Tiere. 284, 1–17 (1965).

116. Ruan J. Bludau S. Palomero-Gallagher N. Caspers S. Mohlberg H. Eickhoff SB. Seitz Rj. Amunts K. Cytoarchitecture, probability maps, and functions of the human supplementary and pre-supplementary motor areas. Brain Struct Funct. 9, 4169–4186 (2018).

117. Bolognini N. Cognitive processes of motor behavior revealed by tDCS. Front Neurol. 8, 29 (2017).

118. Kirchner H. Barbeau EJ. Thorpe SJ. Régis J. Liégeois-Chauvel C. Ultra-rapid sensory responses in the human frontal eye field region. Journal of Neuroscience 29, 7599–7606 (2009).

119. Graziano MSA. The Intelligent Movement Machine. Oxford University Press (2008).

120. Schall JD. On the role of frontal eye field in guiding attention and saccades. Vision Research 44, 1453–1467 (2004).

121. Geyer S. Ledberg A. Schleicher A. Kinomura S. Schormann T. Bürgel U. Klingberg T. Larsson J. Zilles K. Roland PE. Two different areas within the primary motor cortex of man. Nature 382, 805–807 (1996).

122. Geyer S. Schleicher A. Zilles K. Areas 3a, 3b, and 1 of human primary somatosensory cortex. Neuroimage 10, 63–83 (1999).

123. Grefkes C. Geyer S. Schormann T. Roland P. Zilles K. Human somatosensory area 2: observer-independent cytoarchitectonic mapping, interindividual variability, and population map. Neuroimage 14, 617–31 (2001).

124. Scheperjans F. Eickhoff SB. Hömke L. Mohlberg H. Hermann K. Amunts K. Zilles K. Probabilistic Maps, Morphometry, and Variability of Cytoarchitectonic Areas in the Human Superior Parietal Cortex. Cereb. Cortex 18, 2141–2157 (2008).

125. Hülsmann ERM. Traveling cortical netwave, hypothesis. Youtube (2019).

126. Hülsmann ERM. Traveling cortical netwave, results. Youtube (2019).

127. Hülsmann ERM. Traveling cortical netwave, interpretation. Youtube (2019).

